# FiberPro 1.0: Multiagent AI-Guided Design for High-Throughput Production and Conformal Deposition of Functional Protein Micro/Nanofibers

**DOI:** 10.64898/2026.09.18.752775

**Authors:** Longwen Li, Carter Moses, Yechan Noh, Charlie Hsu, Fatima Calderon Gutierrez, Cody Ruiz, Wenjie Liu, Juan Felipe Mogollón Molina, Kaiyu Fu, Huibin Chang

## Abstract

Functional protein micro/nanofibers integrate high specific surface areas with bioactive architectures but remain hampered low production throughput and severe processing instability. Focused rotary jet spinning (FRJS) shows promising to break these throughput constraints while enabling direct, conformal deposition onto complex, irregular substrates. However, navigating FRJS’s narrow processing windows in proteins remains failure-prone without closed-loop experimental guidance. Here, we report FiberPro 1.0, a large language model–driven multi-agent framework that unites high-throughput spinning with real-time experimental feedback for autonomous design, execution, and optimization. Across three protein systems, FiberPro 1.0 converged on spinnable formulations within an average of two iterations. To demonstrate high-throughput conformal deposition in a translational setting, FiberPro designed a zein-based active packaging system applied directly onto diverse food matrices. The resulting conformal coating combined potent antibacterial activity with real-time freshness monitoring, markedly suppressing Escherichia coli and extending shelf life. This work connects autonomous AI reasoning with high-throughput processing, establishing a verifiable route for scalable manufacturing and conformal coating of functional protein micro/nanofibers.

## 1 Introduction

Protein-based micro/nanofibers have emerged as attractive building blocks for functional soft materials due to their high specific surface area, diverse side-chain chemistries, and intrinsic compatibility with bioactive compounds[1,2]. However, existing fabrication techniques, including 3D printing and micro/nanofiber spinning still have unmet needs for the scalable production of protein-based micro/nanofibers. The 3D printing of functional protein micro/nanofibers faces fundamental scalability and processing challenges, especially when the fiber size down to nanometers. Current approaches typically yield sub-gram quantities, far below the throughput required for practical manufacturing. Moreover, conventional 3D printing systems are inherently limited in their ability to deposit micro/nanofibers conformally onto complex, curved, or dynamically moving three-dimensional substrates without sophisticated multi-axis tracking and real-time process[3]. Compared to 3D printing, micro/nanofiber spinning platforms can reach much smaller dimensional features at a relatively higher production rate. Electrospinning is a well-established one for producing protein micro/nanofibers, but its low throughput, high-voltage requirement, solvent dependence, and limited deposition control constrain large-scale manufacturing of functional protein fibers. Harsh processing conditions can also disrupt protein structure and bioactivity. Moreover, electric-field-directed jet trajectories limit precise deposition onto complex or three-dimensional architectures[4].

Rotary jet spinning (RJS) offers a more scalable route to address these limitations[5]. By using centrifugal stretching to draw fibers, RJS can substantially increase production throughput while avoiding the potential perturbation of protein structure and bioactivity[6]. Its further-developed variant, focused rotary jet spinning (FRJS), incorporates a focusing airflow to converge otherwise divergent jets and direct them toward a defined collection region, thereby enabling spatially controlled deposition and three-dimensional fiber assembly[7]. Previous studies have used FRJS to fabricate complex constructs, including heart valves[8], vascular grafts[9], and human-scale heart implants[10]. Nevertheless, the spinnability of protein systems remains far less predictable than that of many synthetic polymers frequently associated with RJS[11]. Protein macromolecules often exhibit insufficient chain entanglement, strong intra- and intermolecular interactions, and pronounced solvent-dependent aggregation, which make empirical optimization slow and unreliable[12,13].

To accelerate materials design, data-driven approaches have recently been introduced[14,15]. Although these methods can facilitate local optimization, they generally face three major limitations: (1) they are often designed to optimize a single target property rather than jointly considering multiple design objectives; (2) they typically require sizable and well curated training datasets, which are rarely available for newly developed protein fiber systems; and (3) they are commonly implemented as open loop recommendation frameworks that cannot integrate heterogeneous scientific evidence, including literature knowledge, characterization images, graphical data and newly generated experimental results, to iteratively refine experimental strategies.

Large language model (LLM)–driven agents offer opportunities to address these challenges, with growing capabilities in scientific reasoning, chemical synthesis planning, and automated research workflows[16–18]. Building on these developments, we introduced FiberPro 1.0 (hereafter, FiberPro), an LLM-based multi-agent system for protein-fiber materials research. In this framework, FiberPro functions as a formulation-discovery and decision-support layer that integrates literature evidence, material constraints, and experimental outcomes, where FRJS serves as the high-efficiency manufacturing platform. A Central Agent coordinates four specialized agents responsible for design, experimentation, analysis, and data management. Together, these agents decompose research tasks, propose candidate strategies, interpret experimental outcomes, and recommend subsequent optimization steps. As an end-to-end demonstration, FiberPro guided the development of an active protein-fiber coating with antibacterial activity and pH-responsive freshness-indicating behavior. This demonstration extends the role of FiberPro beyond knowledge retrieval to experimentally informed, iterative materials development.

To the best of our knowledge, this work presents the first protein fiber research framework that unifies LLM based multi-agent reasoning with micro/nanofiber manufacturing. This framework accelerates the translation of application requirements into manufacturable micro/nanofiber systems and provides a generalizable paradigm for AI guided high throughput and conformal deposition of functional protein micro/nanofibers.

## 2 Results

### 2.1 FRJS as a scalable platform for protein micro/nanofiber fabrication

Despite substantial advances, protein micro/nanofiber fabrication remains limited by process-level bottlenecks, particularly low production throughput, slow experimental feedback, and restricted fabrication of non-planar architectures[19]. To address these challenges, focused rotary jet spinning (FRJS) is used as a scalable manufacturing platform for protein micro/nanofibers. In FRJS, a protein solution is continuously delivered into a high-speed rotating spinneret and expelled via centrifugal force through external orifices. A coaxial focused air stream then simultaneously applies aerodynamic drag and directional confinement and guiding fiber deposition into a defined collection region (Fig.1a). As the jets travel through air, rapid solvent evaporation and centrifugal-force-driven elongation convert the liquid streams into continuous micro/nanofibers.

A key feature of FRJS is the rapid feedback it provides on fiber formation. Nascent fibers appeared within 1 s, and a continuous fiber network formed around the collector within 10 s (Fig. 1b). This response allowed protein spinnability to be assessed within seconds, without prolonged adjustment of electric-field stability or coagulation conditions (Fig. S1). FRJS thus provided a practical platform for rapidly distinguishing nonspinnable formulations from those capable of producing continuous fibers.

**Fig. 1.**
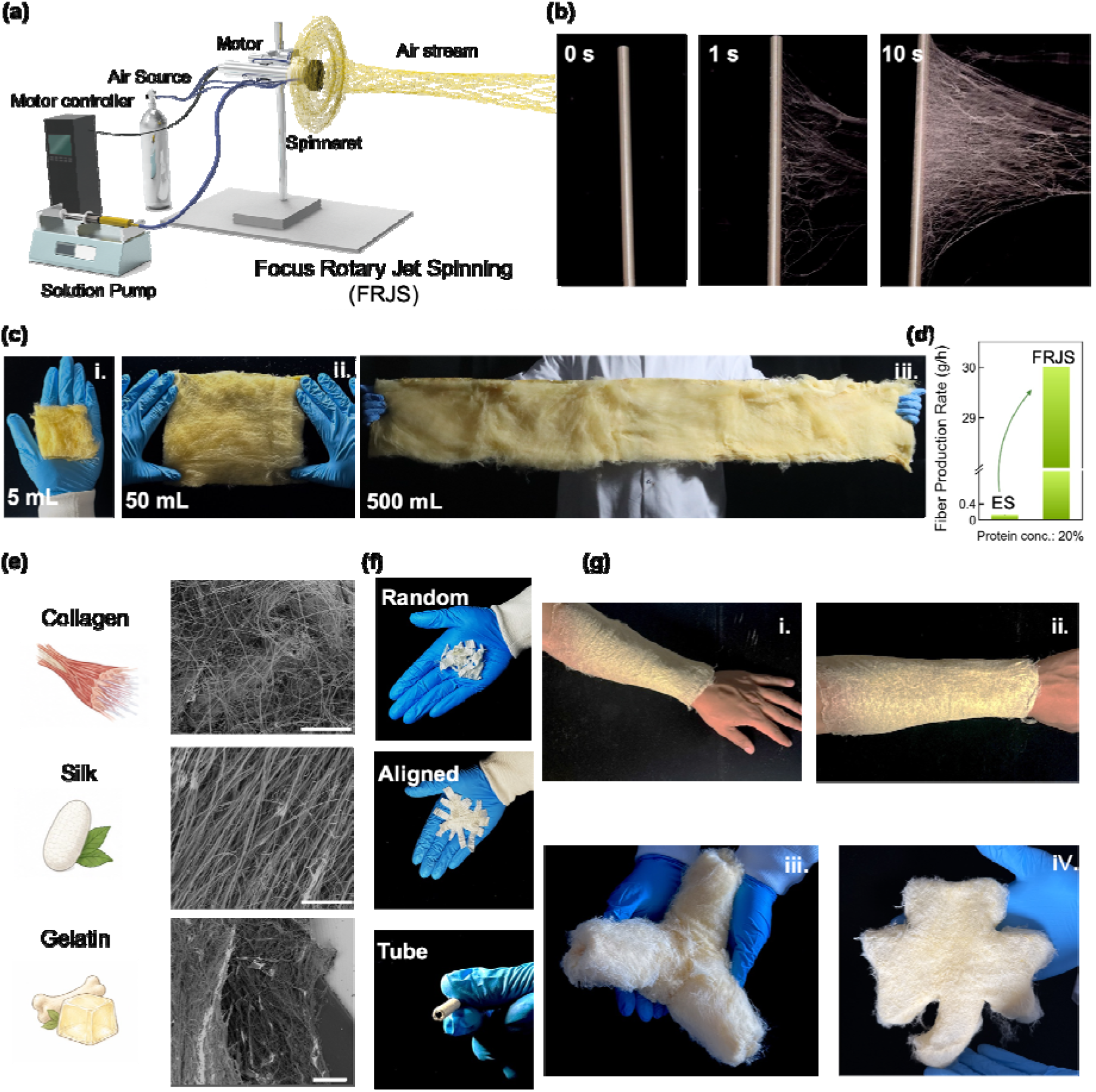
FRJS platform for scalable fabrication of protein micro/nanofibers. (a) Schematic of the focused rotary jet spinning (FRJS) system, comprising a high-speed rotating spinneret, a coaxial focusing air jet, an airflow-guiding shroud, a solution pump, an air source, and a motor controller. (b) Time-lapse images showing rapid fiber formation and continuous deposition during FRJS. (c) Representative zein-fiber mats produced from increasing solution volumes of 5, 50, and 500 mL, illustrating process scalability. (d) Comparison of single-orifice fiber production rates for electrospinning (ES) and FRJS at a protein concentration of 20 wt%. (e) Representative scanning electron microscopy (SEM) images of collagen, silk fibroin, and gelatin fibers fabricated by FRJS. (f) Fiber assemblies with different architectures, including randomly oriented mats, aligned fibers, and tubular constructs. (g) Representative photographs of conformally deposited protein-fiber assemblies, including wearable sleeve-like structures and free-form fibrous constructs. The illustrative icons in panel (e) were created with assistance from ChatGPT (OpenAI) and subsequently reviewed and edited by the authors. These icons are conceptual illustrations and do not represent experimental data.

FRJS also enabled high-throughput production over a range of scales. Increasing the total volume of spinning solution allowed the same setup to transition from small exploratory trials to the continuous production of large-area micro/nanofiber mats (Fig. 1c). A representative mat was first prepared from 5 mL of solution. After verifying the formulation, we increased the processing scale to 50 mL over 20 min and 500 mL over 3 h, producing macroscopic nonwoven protein-fiber sheets. At a typical protein concentration of 20 wt%, FRJS achieved a fiber production rate of approximately 30 g h ¹, nearly two orders of magnitude higher than that of single-needle electrospinning at the same concentration (Fig. 1d and Video S1)[20].

This rapid, high-yield, and programmable manufacturing capability can be generalized across protein systems. We applied FRJS to three protein precursors with distinct biological origins and molecular properties: collagen, silk fibroin, and gelatin (Fig. 1e). Despite marked differences in molecular conformation and aggregation behavior, all three formed continuous micro/nanofiber networks. This breadth establishes FRJS as a general experimental platform for rapidly screening protein fiber formation across diverse protein chemistries.

FRJS also showed clear structural programmability. By varying collector geometry, the same formulation could be processed into random fiber mats, aligned fiber bundles, tubular structures, conformal coatings on human appendages and customized three-dimensional shapes (Fig, 1f, g). Conventional membrane-based nanofiber workflows typically produce planar mats that must subsequently be cut, transferred, rolled, or laminated to obtain the desired geometry. In contrast, FRJS defines the macroscopic shape during deposition, reducing the need for these subsequent shaping and assembly steps.

Together, these capabilities make FRJS well suited to serve as the experimental manufacturing component of an iterative workflow for protein-fiber formulation screening and functional materials development.

### 2.2. AI-Guided Development of Functional Protein Micro/Nanofibers

Although FRJS offers clear advantages for protein micro/nanofiber fabrication, the formulation space for different proteins—including suitable spinning aids and processing windows—remains poorly defined. Changing the protein source or introducing a new functional requirement often necessitates empirical adjustment of the formulation. This trial-and-error process is time-consuming and makes it difficult to consolidate experimental knowledge systematically.

To reduce reliance on experience-driven optimization, we developed FiberPro, an LLM-based multiagent framework that links formulation reasoning to experimental validation and iterative refinement. Rather than replacing researcher judgment, FiberPro supports the formulation, testing, and revision of materials-design hypotheses. The core prompts, representative interactions, and system configurations used in this study are provided in the Supporting Information (File S1)

As shown in Fig. 2a and Fig. S2, FiberPro comprises a (1) Central Agent and four task-specific agents. The Central Agent identifies the material problem raised by the researcher and coordinates the subsequent workflow. (2) Data Agent maintains the literature records, experimental data, structured sample information and uploaded files associated with each project. This shared workspace allows later design decisions to build on accumulated evidence rather than on isolated experiments. (3) Design Agent develops an initial formulation by considering the target property, protein solubility and relevant processing constraints. Once the design is approved, (4) Experiment Agent translates it into an executable wet-lab plan and identifies the variables requiring priority evaluation. After spinning and characterization, (5) Analysis Agent integrates the resulting images, spectra and functional-test data to assess whether the fibers meet the intended criteria. It then recommends the next optimization step.

**Fig. 2.**
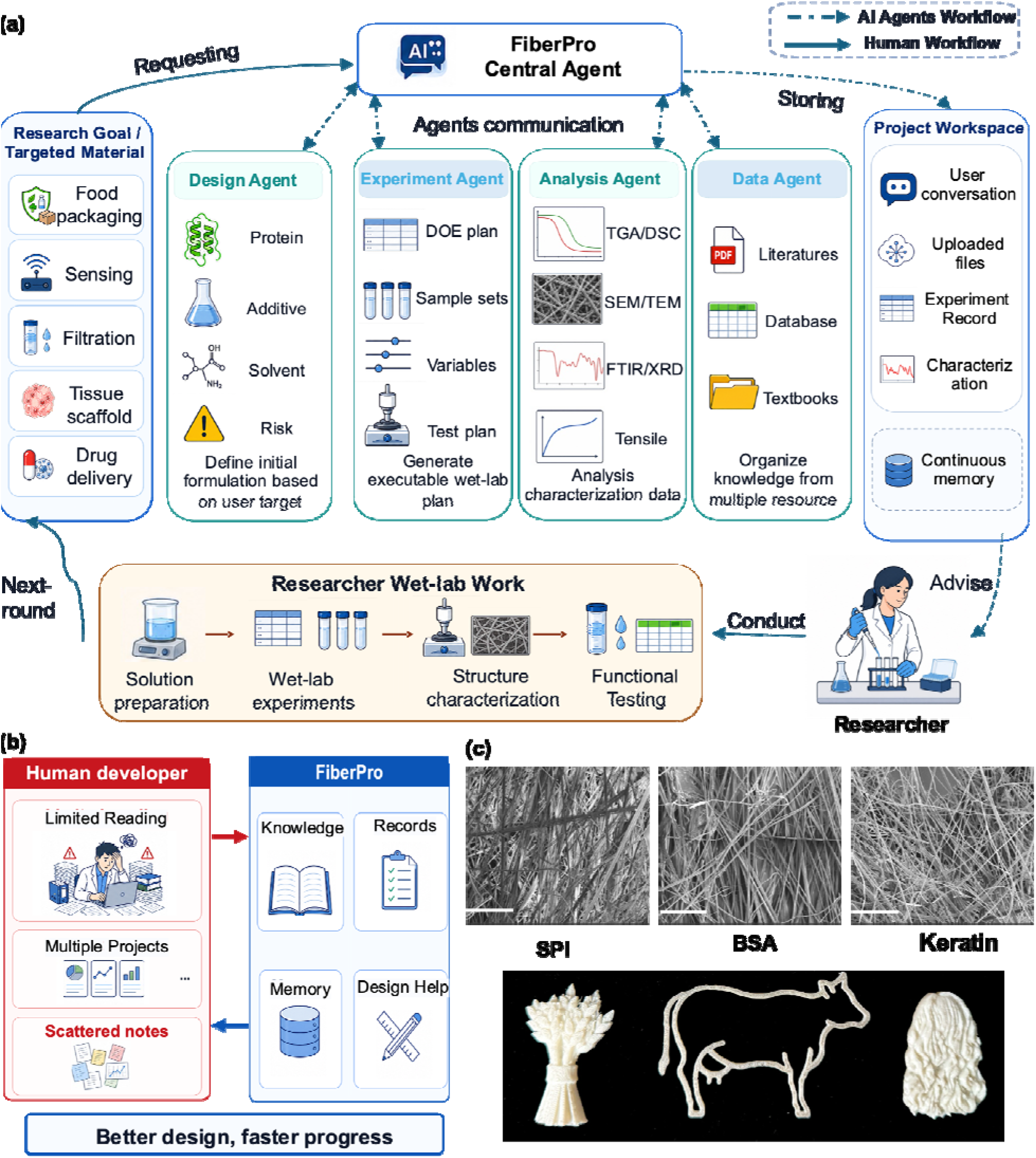
FiberPro-assisted closed-loop development of functional protein micro/nanofibers. (a) Human-in-the-loop workflow of the FiberPro multiagent framework. A researcher specifies a research goal or target material, and the Central Agent coordinates four specialized agents: the Design Agent proposes formulations; the Experiment Agent develops design-of-experiments (DOE) plans; the Analysis Agent interprets characterization and functional testing results; and the Data Agent organizes information from the literature, databases, textbooks, and project records. The agents share a workspace containing user conversations, uploaded files, experimental records, characterization data, and persistent project memory. Human researchers prepare solutions, conduct experiments, characterize materials, and evaluate functional performance, feeding the results back into subsequent design rounds. Dashed and solid arrows denote AI-agent and human workflows, respectively. (b) Comparison of conventional researcher-driven development, in which knowledge and records may be dispersed, with the integrated knowledge, experimental records, project memory, and design support provided by FiberPro. (c) Representative source materials (bottom) and corresponding fibrous products (top) prepared from SPI, BSA, and keratin using FiberPro-guided formulations. Scale bars, 100 μm. The illustrative icons in panels (a) and (b) were created with assistance from ChatGPT (OpenAI) and subsequently reviewed and edited by the authors. These icons are conceptual illustrations and do not represent experimental data.

In a typical workflow (Figs. S3–S7), the researcher first specifies an application goal or materials challenge, such as improving protein spinnability or developing antibacterial protein micro/nanofibers. The Central Agent routes the request to the Design Agent, which refines the objective with the researcher and proposes an initial formulation. Following approval, the Experiment Agent generates a focused design-of-experiments (DOE) plan to evaluate the most informative conditions. The researcher then performs solution preparation, FRJS fabrication, structural characterization, and functional testing. Experimental records are uploaded to the shared workspace, where the Analysis Agent interprets the results and recommends a direction for the next experimental round. This human-in-the-loop process provides an iterative, continuously updatable framework for functional protein micro/nanofiber development (Fig. 2b).

To assess whether FiberPro could extend FRJS to additional protein systems, we selected soy protein isolate (SPI), keratin, and bovine serum albumin (BSA), for which no previous FRJS processing had been reported to our knowledge. Table S1 summarizes representative dialogues, formulation recommendations, and experimental decision pathways for these three systems.

For SPI, FiberPro identified solution homogeneity as the initial constraint. The first design therefore focused on improving dissolution under alkaline conditions. FiberPro subsequently recommended increasing the SPI concentration to raise the effective macromolecular content of the solution. When these adjustments did not yield continuous fibers, FiberPro proposed adding poly(vinyl alcohol) (PVA) as a spinning aid, enabling the formation of a continuous fibrous network.

For keratin, the primary challenge was to maintain solution processability and stable fiber formation at a relatively high protein content. FiberPro proposed a formic acid/poly(vinylpyrrolidone) (PVP) formulation, in which formic acid promoted keratin dissolution and PVP supplied the chain entanglement needed for continuous fiber formation. BSA was used to evaluate the development of an aqueous protein-fiber formulation without conventional organic solvents. FiberPro initially proposed a concentrated aqueous BSA solution containing glycerol as a plasticizer and solution modifier. The first experimental round, however, did not yield collectable fibers. In response, FiberPro identified insufficient chain entanglement and jet stability as the working explanation and recommended gelatin or pullulan as a spinning aid. After the researcher selected pullulan, collectable fibers were obtained in the second experimental round. FiberPro was therefore able to update its materials-design hypothesis in response to failure, rather than conducting unguided repeated screening within a predefined formulation space (Fig.2c, S8).

Overall, FiberPro translated the open-ended question of protein spinnability into a sequence of specific, testable hypotheses and refined its working interpretation as experimental evidence accumulated. For protein systems without an established FRJS processing route, the framework provided an actionable starting point and a traceable decision pathway for subsequent optimization.

### 2.3. AI-guided conversion of zein into active food-coating fibers

To evaluate FiberPro beyond spinnability assessment, we tasked it with developing a functional food-contact coating from a protein with limited precedent in FRJS processing. Zein was selected as the model protein because it is derived from corn and is generally recognized as safe (GRAS) for food-related use [21]. Its intrinsic hydrophobicity makes it attractive for food-contact applications, but obtaining continuous zein fibers by FRJS remained a processing challenge.

As illustrated in Fig. 3a, FiberPro initially selected 70% ethanol to dissolve neat zein while retaining a food-grade formulation (Fig. S9). Across a zein concentration range of 15–35 wt%, however, neat zein formed only particles and short fiber fragments rather than continuous fibers (Fig. S10). FiberPro attributed this failure to insufficient interchain connectivity within zein-rich domains. It therefore recommended poly(ethylene oxide) (PEO) at 5–15 wt% relative to zein as a bridging component to increase network connectivity (Fig. S11). PEO enabled continuous fiber formation, although pronounced local fiber fusion remained (Figs. S12 and S13). After evaluating the SEM images, FiberPro recommended reducing the PEO content and increasing the rotational speed (Fig. S14). These adjustments suppressed fiber fusion and produced continuous fibers with a more uniform morphology.

**Fig. 3.**
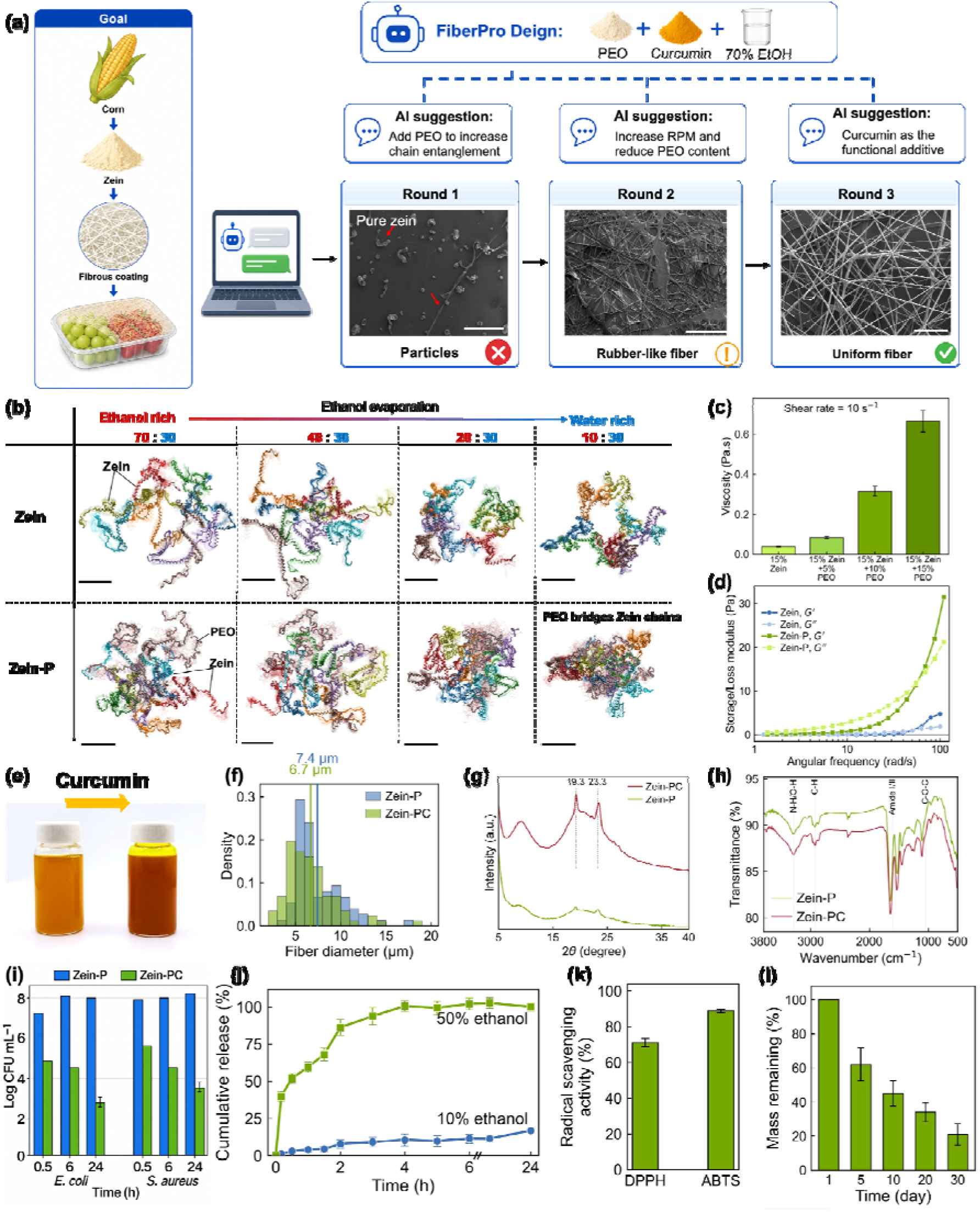
FiberPro-guided closed-loop development and structural and functional characterization of Zein-PC fibers. (a) FiberPro-guided development of zein fibers for food-contact coating applications. Corn-derived zein was formulated in 70% ethanol, with PEO and CUR incorporated during successive design rounds. In round 1, zein alone formed particle-like deposits rather than continuous fibers, prompting a recommendation to add PEO to promote chain entanglement. In round 2, increasing the rotational speed and reducing the PEO content yielded rubber-like fibers with a nonuniform morphology. In round 3, the CUR-containing Zein-PC formulation yielded continuous, interconnected fibers. Representative SEM images show the morphology obtained in each round. Scale bars, 500 μm (rounds 1 and 2) and 200 μm (round 3). (b) Representative equilibrated configurations from all-atom molecular dynamics simulations of zein-only (top) and Zein-P (bottom) systems at ethanol/water volume ratios of 70:30, 40:30, 20:30, and 10:30. Zein chains are represented as differently colored ribbons, and PEO chains are shown using CPK representations (carbon, gray; oxygen, red; hydrogen, white). Translucent replicas show configurations sampled every 0.8 ns over the final 8 ns of each 50 ns trajectory. In the Zein-P system, PEO interacts with multiple zein chains and bridges neighboring chains under the 10:30 condition. (c) Viscosity of spinning solutions containing 15% zein and different PEO contents, measured at a shear rate of 10 s ¹. (d) Frequency-dependent storage modulus (G′) and loss modulus (G″) of neat zein and Zein-P spinning solutions. (e) Photographs of Zein-P and Zein-PC spinning solutions before and after CUR addition, respectively. (f) Fiber-diameter distributions of Zein-P and Zein-PC, with mean diameters of approximately 7.4 and 6.7 μm, respectively. (g) XRD patterns of Zein-P and Zein-PC fibers, with characteristic reflections at 19.3° and 23.3°. (h) FTIR spectra of Zein-P and Zein-PC fibers, with labeled bands corresponding to N–H/O–H, C–H, amide I, and C–O–C vibrations. (i) Antibacterial activity of Zein-P and Zein-PC fibers against *E. coli* and *S. aureus*. (j) Cumulative CUR release from Zein-PC fibers in 10% and 50% aqueous ethanol food simulants. (k) DPPH and ABTS radical-scavenging activities of Zein-PC fibers. (l) Mass remaining after soil burial for 1, 5, 10, 20, and 30 days, showing progressive mass loss. The illustrative icons in panel (a) were created with assistance from ChatGPT (OpenAI) and subsequently reviewed and edited by the authors. These icons are conceptual illustrations and do not represent experimental data.

Rheological measurements supported the proposed role of PEO. Neat zein solutions had low viscosity, whereas PEO incorporation substantially increased solution viscosity. At high angular frequencies, PEO-containing formulations exhibited storage moduli of approximately 20–30 Pa (Fig. 3c, d). This enhanced elastic response is consistent with the network elasticity required for FRJS procesing[22]. All-atom molecular dynamics simulations provided further insight into the contribution of PEO to network formation. Across simulated solvent compositions ranging from ethanol-rich to water-rich conditions, zein chains adopted increasingly compact, aggregated conformations (Fig. 3b). Without PEO, interchain connectivity depended primarily on direct zein–zein contacts. With 10 wt% PEO, individual PEO chains interacted with multiple zein chains, creating additional bridging pathways (Fig. S15). As the ethanol/water volume ratio decreased from 70:30 to 10:30, the zein-chain diffusion coefficient decreased from approximately 0.014 to 0.004 nm² ns ¹. Over the same range, the number of zein–PEO hydrogen bonds per PEO group increased from approximately 0.004–0.007 to approximately 0.014 (Fig. S16). These results support the formation of a more connected transient network, in which strengthened zein–PEO interactions restrict protein-chain mobility and facilitate continuous centrifugal spinning.

FiberPro subsequently selected curcumin (CUR) as a functional additive because of its antioxidant, antibacterial, and pH-responsive properties[23]. CUR addition changed the spinning solution from pale yellow to deep orange (Fig. 3e). The resulting CUR-loaded zein–PEO fibers, denoted Zein-PC, exhibited bright fluorescence and a CUR encapsulation efficiency of approximately 90% (Fig. S17). CUR loading has negligible effect on the fiber diameter or morphology (Fig. 3f and Fig. S18). The XRD patterns of Zein-P and Zein-PC showed similar broad amorphous features, together with weak PEO-related reflections at approximately 19.3° and 23.3° (Fig. 3g). No new sharp reflections were detected after CUR loading, indicating the absence of detectable crystalline CUR domains in the fiber matrix. In the FTIR spectra, the N–H/O–H band shifted from 3295 to 3283 cm ¹, the amide I band from 1652 to 1647 cm ¹, and the amide II band from 1538 to 1543 cm ¹ (Fig. 3h)[24]. These shifts are consistent with intermolecular hydrogen-bonding interactions among zein, PEO, and CUR.

We next evaluated the functional performance of the FRJS-fabricated Zein-PC fibers. After 24 h of treatment with Zein-PC fibers, viable counts of *E. coli* and *S. aureus* decreased to approximately 2.8 and 3.5 log CFU mL ¹, respectively (Fig. 3i), demonstrating antibacterial activity against both strains. CUR release from Zein-PC fibers depended strongly on the food simulant. In 50% ethanol, representing an oil-rich food environment, the cumulative CUR release reached approximately 90–95%. In the aqueous food simulant, cumulative release remained below 20% after 24 h (Fig. 3j). This difference is consistent with the hydrophobicity of CUR and its medium-dependent solubility and partitioning behavior. CUR can diffuse more readily from the fiber matrix in ethanol-rich media, whereas its limited aqueous solubility suppresses rapid migration and prolongs retention within the fibers.

Zein-PC fibers also exhibited antioxidant activity, with DPPH and ABTS radical-scavenging activities of approximately 70% and 85–90%, respectively, exceeding those of the unloaded control fibers (Fig. 3k). During soil burial, the fibers progressively lost mass over 30 days, retaining approximately 15–20% of their initial mass at the final time point (Fig. 3l and Fig. S19). Collectively, these results show that FiberPro converted an initially unspinnable zein formulation into a continuous, CUR-loaded food-contact fiber coating with antibacterial, antioxidant and degradable characteristics.

### 2.4. In Situ Conformal Coating of Real Produce for Active Preservation and Intelligent Freshness Prediction

Unlike coating and packaging approaches that require immersion, heat sealing, or adhesives, FRJS enables noncontact conformal fiber deposition directly onto food surfaces without external pressure or post-deposition bonding. Using an apple as a representative substrate, Zein-PC fibers solidified during flight and formed a thin, continuous yellow coating that followed both the overall curvature and local features of the peel (Fig. 4a). The coating could be removed by gentle rinsing with water, leaving no visible residue or surface damage (Video S3). The same deposition process was applied to cherry tomatoes, strawberries, apples, mangoes, and watermelon, demonstrating compatibility with substrates of different sizes and geometries (Fig. 4b–d).

**Fig. 4.**
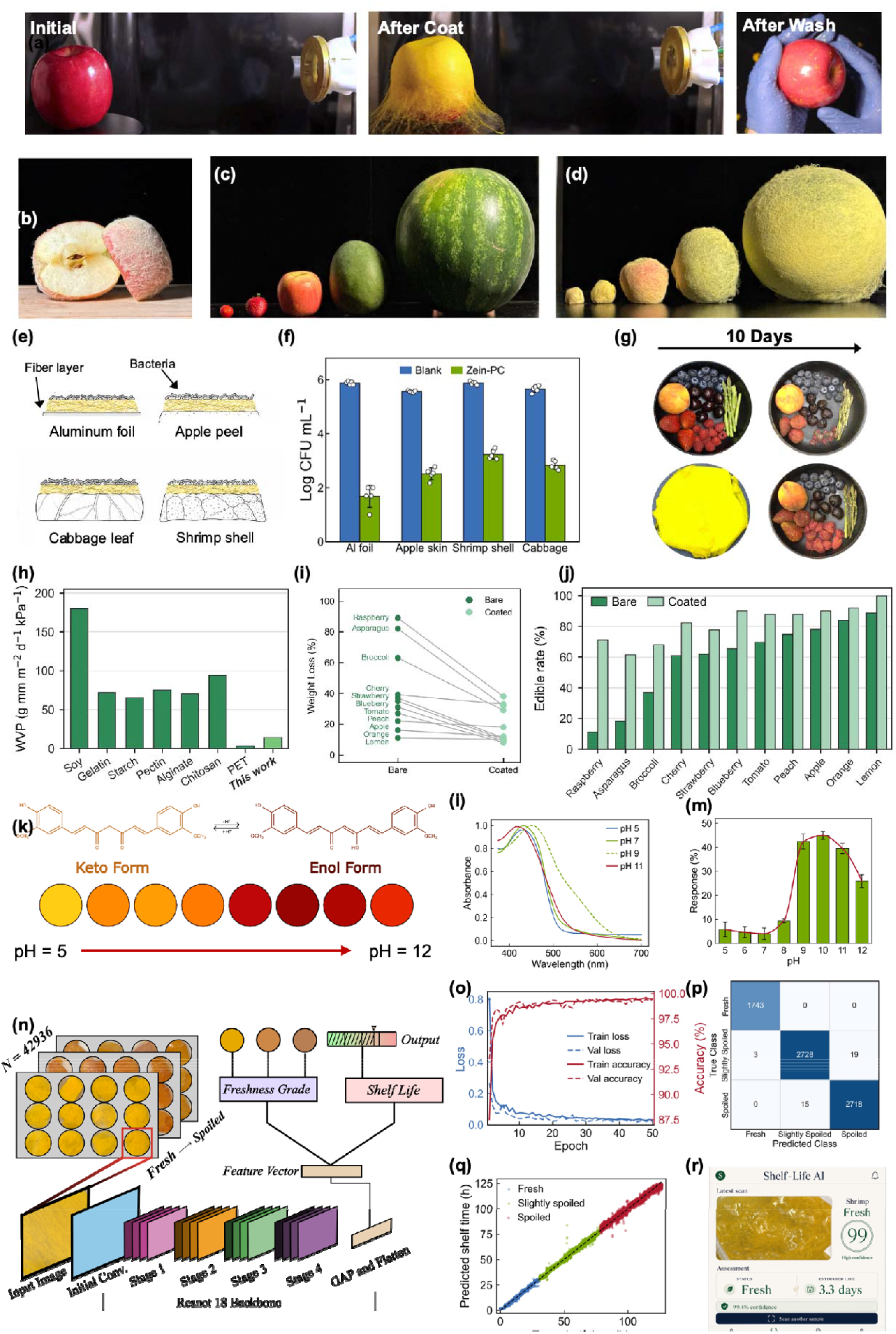
Universal in situ Zein-PC fiber coating for fresh-produce preservation, colorimetric sensing and AI-enabled shelf-life prediction. (a) In situ FRJS deposition of curcumin-loaded zein–PEO (Zein-PC) fibers onto an apple, shown before coating, immediately after deposition and after washing. (b) Cross-sectional photograph of the coated apple, showing conformal coverage of the peel by the fibrous layer. (c, d) Representative fruits and vegetables with varied sizes and geometries before and after FRJS coating, demonstrating the broad substrate compatibility of the deposition process. (e) Schematic of the food-associated surfaces used to evaluate contact-active antibacterial activity, including aluminum foil, apple peel, cabbage leaf and shrimp shell. (f) E. coli counts on the indicated substrates after 24 h of contact with blank or Zein-PC-coated surfaces. (g) Representative photographs of mixed produce before and after 10 d of ambient storage, comparing uncoated (top) and fiber-coated (bottom) samples. (h) Water-vapor permeability (WVP) of Zein–PC fibrous membranes compared with representative packaging materials. (i) Weight loss of different produce items after 10 d of storage with and without fiber coating. (j) Edible rate of individual produce items after 10 d of ambient storage. (k) Keto–enol transformation of curcumin and the corresponding color evolution of Zein-PC fibers from pH 5 to 12. (l) UV–Vis absorption spectra of Zein-PC fibers at selected pH values. (m) Quantified colorimetric response of Zein-PC fibers as a function of pH. (n) Architecture of the multitask convolutional neural network with a ResNet-18 backbone for simultaneous freshness classification and shelf-life regression from coating images. (o) Training and validation loss and classification accuracy during model optimization. (p) Confusion matrix for three-class freshness classification on the held-out test set. (q) Predicted versus measured shelf time for all test samples, colored by freshness class. (r) Shelf-Life AI smartphone application for real-time freshness assessment and shelf-life prediction using the halochromic Zein-PC coating.

To determine whether antibacterial activity was maintained at food interfaces, we evaluated Zein-PC coatings against E. coli on food-relevant substrates (Fig. 4e, f). Aluminum foil served as an inert reference, while apple peel, cabbage leaves, and shrimp shells represented biological surfaces with different compositions and structures. On aluminum foil, Zein-PC reduced bacterial counts from approximately 5.9 to 1.7 log CFU mL ¹. Differences in coating coverage, CUR release, and bacterial attachment may contribute to the substrate-dependent activity observed on biological surfaces. Nevertheless, the coatings achieved reductions of at least approximately 2.7 log units on all tested food interfaces, demonstrating antibacterial activity beyond the inert reference surface.

We next evaluated preservation across 11 perishable foods, including berries, other fruits, vegetables and citrus. After 10 days of storage at room temperature, uncoated samples generally showed water loss, shrinkage, wrinkling, browning and tissue softening, while berries also developed visible mold. In contrast, coated samples retained greater gloss, fullness and structural integrity (Fig. 4g). These outcomes were associated with the transport properties of the Zein-PC coating, which showed a water-vapor transmission rate of approximately 14 g mm m ² d ¹kPa ¹ (Fig. 4h). Compared with gelatin, starch and other biopolymer films, Zein-PC showed lower water-vapor transmission [25]. This balance limited excessive moisture loss while maintaining gas exchange compatible with fresh-produce respiration. Consistently, Zein-PC reduced storage-associated mass loss, particularly for dehydration-prone samples (Fig. 4i), and improved the edible rate of all tested foods after 10 d (Fig. 4j).

Beyond active preservation, Zein-PC retained the pH-responsive colorimetric behavior of curcumin (CUR). As pH increased from 5 to 12, shifts in the keto–enol equilibrium of CUR changed the fiber color from yellow and orange yellow to dark red (Fig. 4k). Corresponding UV– Vis spectra showed marked changes in the absorption bands, consistent with altered electronic structure of the CUR chromophore under alkaline conditions (Fig. 4l). The colorimetric response remained at approximately 4–6% between pH 5 and 7, then increased rapidly to approximately 39–45% between pH 9 and 11, reaching a maximum near pH 10 (Fig. 4m and Fig. S20). This response window encompasses the weakly alkaline conditions generated by volatile-amine accumulation during protein-rich food spoilage, providing a material basis for visual spoilage recognition.

Because a single-color measurement cannot capture subtle changes at the onset of spoilage, we next used time-lapse imaging to quantify the rates of color change and surface-morphology evolution. Shrimp, which spoils rapidly and exhibits well-defined changes in biogenic amines, were used as the model food. Under constant 21 °C storage and 5600 K illumination, 42,936 images were collected, capturing the full transition from fresh to mildly spoiled and spoiled states (Fig. 4n and Fig. S21).

Using this dataset, we developed a multitask convolutional neural network with a shared ResNet-18 backbone for simultaneous freshness classification and storage-time regression (Fig. 4n). The shared network extracted multiscale information from color distributions, texture and surface morphology, and two independent output branches returned three freshness classes and continuous storage-time estimates. Training and validation losses decreased rapidly and converged, while classification accuracy stabilized near 100% (Fig. 4o). In an independent test set of 7,226 images, the model correctly classified 7,189 images, yielding an overall accuracy of 99.49% (Fig. 4p). All fresh samples were correctly classified, and most misclassifications occurred between the adjacent mildly spoiled and spoiled classes. Predicted storage times closely matched the measured values, with R² = 0.997, a mean absolute error of 1.40 h, and a root-mean-square error of 1.78 h (Fig. 4q). The predicted median storage times for the three classes were 14.45, 55.60, and 103.12 h, compared with measured medians of 15.01, 54.82, and 102.40 h, respectively. The approximately 41–48 h separation between adjacent stages exceeded the prediction error (Fig. S22). These results show that coating images captured distinguishable stages along the spoilage trajectory under the tested conditions, providing a basis for image-assisted freshness assessment and sorting.

To translate the material response and image analysis into an accessible interface, we integrated the prediction workflow into a lightweight prototype smartphone application, Shelf-Life AI (Fig. 4r). Images of Zein-PC-coated foods acquired with a conventional smartphone camera returned a freshness class, an estimated remaining shelf life, and model confidence within approximately 2–3 s. The prototype provides a camera-based interface for material-assisted freshness assessment.

Together, the FiberPro-designed Zein-PC system integrates in situ fiber deposition, interfacial antibacterial activity, moisture-loss regulation, colorimetric sensing and intelligent freshness prediction within a single food-coating platform. It thereby establishes a continuous application pathway from material design to validation on real foods and mobile decision support.

## 3. Discussion

This study establishes an experimentally guided framework for protein micro/nanofiber development by integrating FRJS with FiberPro. FRJS provides rapid, high-throughput fabrication and control over the architecture of deposited fibers, while FiberPro serves as a human-in-the-loop, LLM-based decision-support layer linking material constraints, formulation design, experimental planning, characterization, and subsequent optimization. Together, these components organize protein-fiber development into a traceable sequence of hypotheses, experiments, and evidence-based revisions rather than isolated rounds of empirical screening.

Across soy protein isolate, keratin and bovine serum albumin systems lacking prior FRJS precedent, FiberPro proposed distinct routes to address material-specific limitations, including improved dissolution, increased effective macromolecular content, enhanced chain entanglement and improved jet stability. These results show that the value of FiberPro does not lie in applying a single rule to predict the spinnability of all proteins. Rather, it updates material-design hypotheses in response to experimental failures and provides targeted starting points for subsequent experiments. FiberPro therefore does not replace researcher judgment, but improves knowledge integration, decision recording and iterative optimization during protein-fiber development.

The zein study serves as a representative case demonstrating that this framework can extend beyond fiber formation toward functional application. FiberPro converted an initially unspinnable zein formulation into multifunctional Zein-PC fibers and linked material design with in situ food preservation, colorimetric sensing and image-based shelf-life prediction. This result indicates that the framework can support both the resolution of fundamental processing challenges and the translation of protein-fiber materials toward application-oriented performance.

The present study remains limited to a selected set of protein systems and controlled application settings. The transferability of FiberPro across broader protein classes, its systematic comparison with conventional optimization strategies and its robustness across laboratories require further evaluation. Nevertheless, these findings show that coupling large language model– guided reasoning with rapid fiber manufacturing and experimental feedback provides a scalable human–AI approach for the discovery, optimization and application of functional protein micro/nanofibers.

In summary, coupling LLM-guided formulation reasoning with rapid FRJS fabrication creates an experimentally grounded, human–AI workflow for functional protein micro/nanofiber development. By connecting iterative formulation design, high-throughput production, conformal deposition, and application testing, FiberPro provides a foundation for developing protein-fiber materials tailored to specific functional requirements.

## 4 Experimental Section

### 4.1. Materials

Zein, soy protein isolate, gelatin, collagen, bovine serum albumin (BSA), curcumin, 200-proof ethanol, 2,2-diphenyl-1-picrylhydrazyl (DPPH), 2,2′-azino-bis(3-ethylbenzothiazoline-6-sulfonic acid) (ABTS), poly(ethylene oxide) (PEO; Mw = 60 kDa), poly(vinylpyrrolidone) K90 (PVP), reduced iron powder, and potassium persulfate were purchased from Sigma-Aldrich (St. Louis, MO, USA).

Silk fibroin was extracted from Bombyx mori cocoons by degumming in 0.02 M Na CO, dissolving in 9.3 M LiBr at 60 °C, dialyzing against deionized water, and freeze-drying. Keratin was extracted from wool using a solution containing 8 M urea and 0.14 M dithiothreitol (DTT; pH 10.5) at 90 °C for 4 h, followed by dialysis against deionized water and freeze-drying. Fresh produce was purchased from a local market on the day of each experiment.

### 4.2. Fabrication of protein Micro/nano fibers based on FRJS

Protein micro/nanofibers were fabricated using a custom system comprising a high-speed motor spindle (Model 5000, RpmTech, USA), a stainless-steel spinneret with a 600 μm orifice, a syringe pump (Harvard Apparatus 703007, USA), and a 3D-printed airflow-guiding shroud for jet stabilization. Unless otherwise specified, spinning solutions were delivered at 2.5 mL min ¹ and spun at 8,000 rpm with a spinneret-to-collector distance of 22 cm.[10]

Zein spinning solutions containing 15–35 wt% zein were prepared in 70% (v/v) aqueous ethanol. For the optimized formulation, PEO was added at 10 wt% relative to zein, followed by CUR at 5 wt% relative to zein, to obtain the Zein-PC spinning solution Gelatin, collagen, and silk fibroin spinning formulations were prepared separately in hexafluoroisopropanol (HFIP) at a total polymer concentration of 7 wt%, with polycaprolactone (PCL) incorporated at 30 wt% of the total polymer content as a spinning aid. The aqueous BSA formulation contained 20 wt% BSA, 10 wt% pullulan, and 2 wt% glycerol. Keratin formulations were prepared by dissolving 12 wt% keratin in 90% formic acid and adding 2 wt% PVP K90 as a spinning aid. SPI formulations were prepared by dissolving 15 wt% SPI in alkaline water at pH 11 and adding 5 wt% PVA as a spinning aid. For each protein system, solution composition and spinning parameters were experimentally optimized to obtain continuous fibers. FiberPro-generated formulation recommendations were experimentally validated using FRJS.

### 4.3. Characterizations

#### 4.3.1. Morphological Characterization

Fiber surface morphology was examined using environmental scanning electron microscopy (Prisma Environmental SEM, Thermo Scientific, USA). SEM images were analyzed quantitatively using ImageJ (NIH, USA).

#### 4.3.2. Chemical Structure Characterization

FTIR spectra were acquired using a Nicolet iS50 spectrometer (Thermo Fisher Scientific, USA) in attenuated total reflectance (ATR) mode over 4000–400 cm ¹, with a resolution of 4 cm ¹ and 32 co-added scans.

#### 4.3.3. Rheological Properties Analysis

Rheological measurements were performed at 25 °C using an HR-2 Discovery Hybrid Rheometer (TA Instruments, USA) with a cone-and-plate geometry. Steady-shear viscosity was measured over a shear-rate range of 0.1–1000 s ¹. Oscillatory frequency sweeps were subsequently conducted over 0.1–100 rad s ¹ at a strain amplitude within the linear viscoelastic region to determine the storage modulus (G′) and loss modulus (G″)

#### 4.3.4. X-Ray Diffraction Analysis

X-ray diffraction (XRD) analysis was performed using a SmartLab X-ray diffractometer (Rigaku, Japan). Fiber samples were mounted on a low-background sample holder and scanned in the 2θ range of 5–40° using Cu Kα radiation. Diffraction patterns were collected to evaluate the crystalline and amorphous structures of the protein-based fibers and to compare structural changes induced by different protein formulations and assisting polymers. The obtained XRD profiles were analyzed based on the positions, intensities, and broadening of characteristic diffraction peaks.

#### 4.3.5. Encapsulation Efficiency and In Vitro Release Behavior

To determine the encapsulation efficiency (EE) of the zein micro/nanofibers, a certain amount of the sample was fully dissolved. The curcumin concentration was then determined using a previously established calibration curve (R² > 0.99). EE was calculated according to Equation (1):

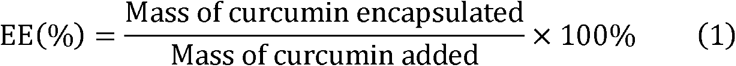

The release behavior of zein-cur micro/nanofibers was assessed using two food simulants (A (10% v/v ethanol) and D1 (50% v/v ethanol)) in accordance with regulatory guidelines set by the U.S. FDA[26]. Curcumin-loaded micro/nanofibers were fully immersed in 20 mL of the simulant and incubated in a shaking water bath at 37 °C. At designated time points, aliquots of the release medium were collected, and curcumin concentration was quantified spectrophotometrically at 425 nm. See Equation (2):

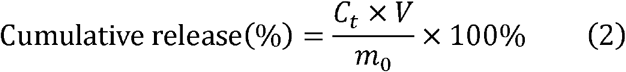

*C_t_* is the curcumin concentration, and *m_0_*is the initial amount.

#### 4.3.6. Water Vapor Permeability

WVP was measured gravimetrically using a standard cup method. Fiber mats were sealed onto cups (10 mm inner diameter) containing 10.0 mL of deionized water and stored at 25 °C, 75% RH. Mass loss was recorded at 24 h intervals over 5 days. WVP was calculated as:

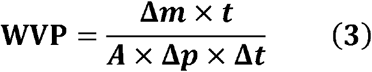

where Δm is the mass loss (g), t is the mat thickness (mm), A is the exposed area (m²), Δp is the water vapor pressure difference (Pa), and Δt is the measurement duration (day).

#### 4.3.7. Antioxidant Activity Assessment

The antioxidant activity of the zein-cur micro/nanofiber samples was assessed using the 2,2-diphenyl-1-picrylhydrazyl (DPPH) radical-scavenging assay. Each micro/nanofiber sample (film-equivalent concentrations of 1–5 mg mL ¹) was combined with 4 mL of a 100 μM DPPH ethanolic solution and incubated in the dark for 1 h at room temperature. Following incubation, the absorbance of the mixture was recorded at 517 nm. The DPPH scavenging activity (%) was then calculated according to Equation (4):

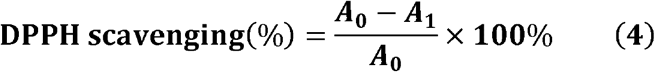

where *A_0_* and *A_1_* represent the absorbance values of the blank and the reaction mixture at 517 nm.

To assess the ABTS• radical-scavenging activity of the micro/nanofiber samples, the ABTS• solution was diluted to reach an absorbance of 0.70 ± 0.02 at 734 nm. Micro/nanofiber samples were dispersed in 10 mL of water and agitated for 1 h. Sample solutions with concentrations of 1.0 - 5.0 mg·mL ¹ were prepared by mixing 1 mL of each supernatant with 3 mL of the ABTS• solution. Samples were then incubated in the dark for 1 h. Absorbance was recorded at 734 nm, and the ABTS• scavenging activity (%) was calculated according to Equation (5):

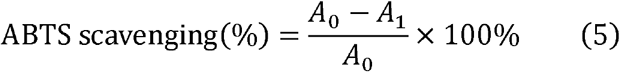

where *A_0_* and *A_1_* denote the absorbance values of the blank and the sample solutions at 734 nm.

#### 4.3.8. Antibacterial Activity Assessment

The antibacterial activity of Zein-PC composite micro/nanofibres was evaluated using suspension-based and surface-based direct-contact assays. Suspension assays were performed against *Escherichia coli* (ATCC 25922) and *Staphylococcus aureus* (ATCC 25923). For both strains, Zein-P and Zein-PC fiber mats were compared. Circular specimens (2 cm diameter) were UV-C sterilized and incubated in inoculated, strain-appropriate growth medium at 37 °C. At 0.5, 6 and 24 h, samples were serially diluted, plated on the corresponding agar medium and quantified by colony-forming unit (CFU) enumeration. Antibacterial activity was expressed as viable counts and log reductions relative to the PBS-treated control.

Surface-based direct-contact activity was evaluated against E. coli on food-relevant substrates. Apple peels, cabbage leaves, and shrimp shells were cut into 2 cm × 2 cm specimens and coated in situ with Zein-PC micro/nanofibers by FRJS. Uncoated specimens served as substrate-matched controls, and aluminum foil was included as an inert reference. An E. coli-containing agar slurry comprising 0.85% NaCl, 0.3% agar, and 1 × 10 –3 × 10 CFU mL ¹ was applied to each specimen at a volume of 300 μL, corresponding to an initial load of approximately 3 × 10 –9 × 10 CFU per specimen. After gelation for 5 min, samples were analyzed immediately or incubated for 24 h at 37 °C and approximately 80% relative humidity. Surviving bacteria were recovered using sterile PBS and quantified by CFU enumeration. Results were expressed as viable counts and log reductions relative to the corresponding control. Each condition was tested in three independent replicates

#### 4.3.9. Produce quality and shelf-life assessment

Shelf-life performance was evaluated using representative perishable foods, including strawberry, cherry, raspberry, blueberry, peach, asparagus, broccoli, lemon, tomato, orange, and apple. After non-contact, in situ deposition of zein–PEO–CUR micro/nanofibers by FRJS, coated samples and uncoated controls were stored under ambient conditions (20–25 °C, 50–60% relative humidity) for up to 10 days. Samples were visually inspected and imaged at defined time points. Spoilage rate was defined as the fraction of spoiled samples at each time point. Weight loss was measured as an indicator of dehydration by recording sample mass at Day 0. The visually assessed edible fraction was defined as the mass fraction of tissue without visible browning, mold, or structural collapse

### 4.4 Colorimetric Analysis

#### 4.4.1 Colorimetric Response of Zein–PEO–CUR Fibers

Zein-PC fiber mats were immersed for 20 min in phosphate–borate–citrate buffers spanning pH 5–12. Images were acquired in a light-controlled chamber under 5600 K LED illumination using a Canon EOS 90D camera equipped with a 50 mm f/1.8 lens. ISO, aperture, white balance, and working distance were held constant. A custom Python script was used to segment the fiber regions, convert the images to CIELAB color space, and calculate mean L*, a*, and b* values.

#### 4.4.2 Machine Learning Model and App Development

Shrimp samples coated with Zein-PC fibers were imaged at 5 min intervals under the illumination conditions described in Section 4.4.1, yielding 42,936 images. The dataset was partitioned by individual shrimp specimen into training, validation, and held-out test subsets at a ratio of 70:15:15 to prevent images of the same specimen from appearing in different subsets. Images were resized to 224 × 224 pixels, and data augmentation included random horizontal flips, rotations of up to ±15°, and brightness jitter. A multitask convolutional neural network (MT-CNN) was constructed using a pretrained ResNet-18 backbone with shared parameters and two task-specific heads. The classification head predicted fresh, mildly spoiled, or spoiled states using cross-entropy loss. The regression head predicted storage time in hours using mean squared error loss, weighted by λ = 0.001. The model was trained using the Adam optimizer with a learning rate of 0.001 and a batch size of 32 for up to 50 epochs, with early stopping based on validation loss. Model training was performed in PyTorch on an NVIDIA RTX A1000 GPU. The trained model was exported to TensorFlow Lite and deployed in a prototype smartphone application for on-device freshness inference.

### 4.5 All-atom Molecular Dynamics Simulations of Solvent-dependent Zein–PEO Aggregation, Connectivity and Chain Mobility

All-atom molecular dynamics (MD) simulations examined how ethanol ratio and poly(ethylene oxide) (PEO) affect α-zein aggregation, interchain packing and mobility. α-Zein Z19 (UniProt P06677, Zea mays) was modeled as its mature 219-residue sequence (residues 22– 240; signal peptide removed). Because the AlphaFold [27] structure is largely disordered and has low confidence (global pLDDT ≈ 47), one chain was first collapsed by brief equilibration in ethanol/water solvent. This compact conformer seeded the multi-chain systems, allowing aggregation to emerge rather than be imposed. Hydroxyl-terminated PEO 109-mers (4.8 kDa) were built with the CHARMM-GUI Polymer Builder [28,29]. Zein, ethanol, PEO and water were described by CHARMM36m [30], CGenFF [31], refined CHARMM36 poly(ethylene glycol)/ether parameters [32], and CHARMM-modified TIP3P [33,34], respectively.

Each system contained eight zein chains, with or without four PEO chains at 10 wt% of the zein mass. Simulations used ethanol volume ratios of 70:30, 40:30, 20:30 and 10:30 at 20 g zein per 100 mL solvent. Solvent molecule counts were set using the target volume ratio and volume-weighted solvent density, for example, 9,430 ethanol and ∼22,850 water molecules at 40:30. Chains were packed into a ∼13.4 nm cubic box with GROMACS insert-molecules, then solvated with ethanol and water. Cl− counterions neutralized the net charge, giving ∼180,000 atoms per system.

Following steepest-descent minimization, systems were equilibrated for 0.3 ns in NVT and 20 ns in NPT at 300 K. The C-rescale barostat [35] was applied isotropically (τp = 2 ps, 10 bar) to establish fluid density. Systems were then annealed for 50 ns in NVT at 400 K, followed by 5 ns of NVT cooling to 300 K. The elevated-temperature stage was run at fixed volume to promote mixing and interchain contact while avoiding ethanol vaporization. Hydrogen mass repartitioning (factor 3) [36] enabled a 4 fs timestep during annealing, cooling and production; initial equilibration used a 2 fs timestep. LINCS [37] constrained all bonds involving hydrogen. Temperature was controlled with the v-rescale thermostat [38] (τT = 0.1 ps), electrostatics with particle-mesh Ewald [39], and van der Waals interactions with a 1.0–1.2 nm force-switch.

Production simulations were run for 50 ns in NVT at 300 K, with three independent replicates per system and coordinates saved every 40 ps. Paired zein-only and zein+PEO systems shared identical initial zein coordinates within each replicate. Simulations used GROMACS 2025.2 [40] on NVIDIA A40 GPUs (NCSA Delta).

Properties were analyzed over the final 30 ns. Intermolecular hydrogen bonds were counted using donor–acceptor distances ≤ 3.5 Å and D–H···A angles ≥ 150°. Aggregate size and chain mobility were quantified by the heavy-atom radius of gyration after periodic-boundary reassembly and heavy-atom mean-squared displacement, respectively. Interchain association was represented as a contact graph of the eight zein chains, with edges assigned to pairs exceeding a threshold number of close contacts and PEO-mediated bridges added in zein+PEO systems. Values are mean ± SEM across three replicates. Analyses used in-house code based on MD Analysis [41,42].

## Supporting information

Support Information

## Acknowledgements

This work was supported by start-up funding from the University of Notre Dame. The authors acknowledge the Materials Characterization Facility (MCF) at the University of Notre Dame for access to the rheometer and Fourier transform infrared (FTIR) spectroscopy instrumentation. The MCF is supported by Notre Dame Research. This work used the Delta GPU system at the National Center for Supercomputing Applications (NCSA) through allocation PHY250014 from the Advanced Cyberinfrastructure Coordination Ecosystem: Services & Support (ACCESS) program (Award No. PHY250014), which is supported by U.S. National Science Foundation grants #2138259, #2138286, #2138307, #2137603, and #2138296. This work was supported by the research fund provided to Y.N. through the University of Notre Dame Bioengineering and Life Sciences (BELS) Postdoctoral Fellowship.

## CRediT authorship contribution statement

**Longwen Li**: Conceptualization, Methodology, Investigation, Formal analysis, Writing – review & editing. **Carter Moses**: Conceptualization, Methodology, Investigation, Formal analysis, Writing – review & editing.**Yechan Noh**: Investigation, Formal analysis, Writing – review & editing. **Charlie Hsu**: Investigation, Writing – review & editing. **Fatima Calderon Gutierrez**: Investigation, Validation, Writing – review & editing. **Cody Ruiz**: Resuorces, Validation, Writing – review & editing. **Wenjie Liu**: Writing – review & editing. Juan **Felipe Mogollón Molina**: Investigation, Writing – review & editing. **Kaiyu Fu**: Supervision, Writing – review & editing. **Huibin Chang**: Resuorces, Supervision, Writing – review & editing.

## Data availability

The data that support the findings of this study are available from the corresponding author upon reasonable request.

## Declaration of generative AI in the manuscript preparation process

During the preparation of this manuscript, the authors used ChatGPT (OpenAI) to assist with the creation of illustrative icons in Figs. 1e, 2a–b, and 3a. These elements were used solely for conceptual illustration and do not represent experimental observations or data. The authors reviewed and edited the generated content and take full responsibility for the published article.

