## Supplementary material for "FiberPro 1.0: Multiagent AI-Guided Design for High-Throughput Production and Conformal Deposition of Functional Protein Micro/Nanofibers": Support Information

Longwen Li *et al.*

**This PDF file includes:**

Figs. S1 to S22

Table S1

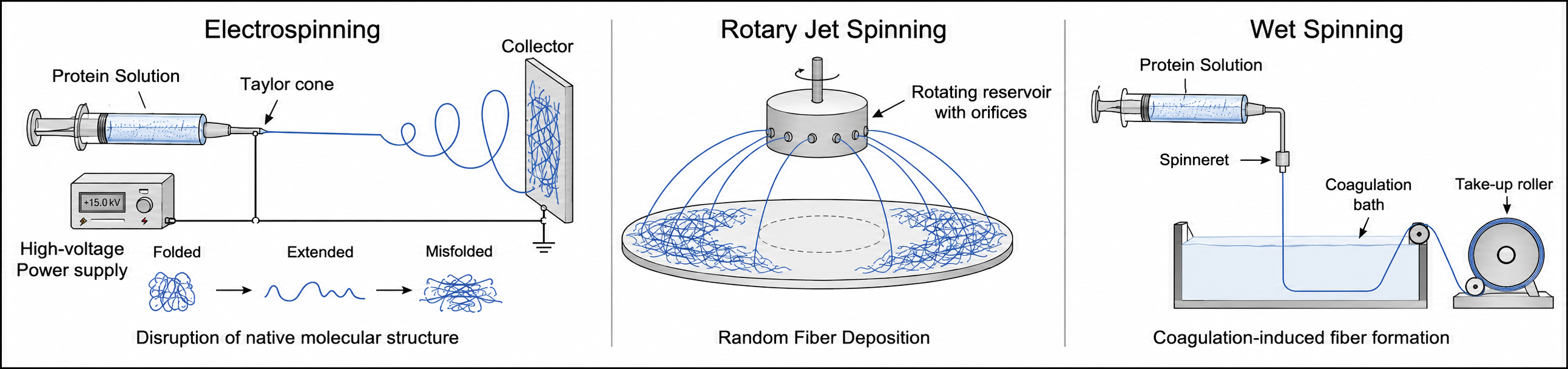

**Fig. S1. Comparison of conventional protein fiber fabrication methods and their limitations.** Electrospinning uses a high-voltage electric field to draw a protein solution into fine fibers, but its relatively low throughput and strong electric field may perturb the native structure of sensitive proteins. Conventional RJS offers substantially higher fiber production rates than electrospinning but provides limited control over fiber trajectories and deposition, complicating the fabrication of spatially defined or conformal fibrous architectures. Wet spinning extrudes a protein solution into a coagulation bath, where solvent exchange or nonsolvent-induced phase separation promotes continuous fiber formation; however, the process requires carefully controlled coagulation conditions and is primarily suited to the production of continuous, one-dimensional fibers.

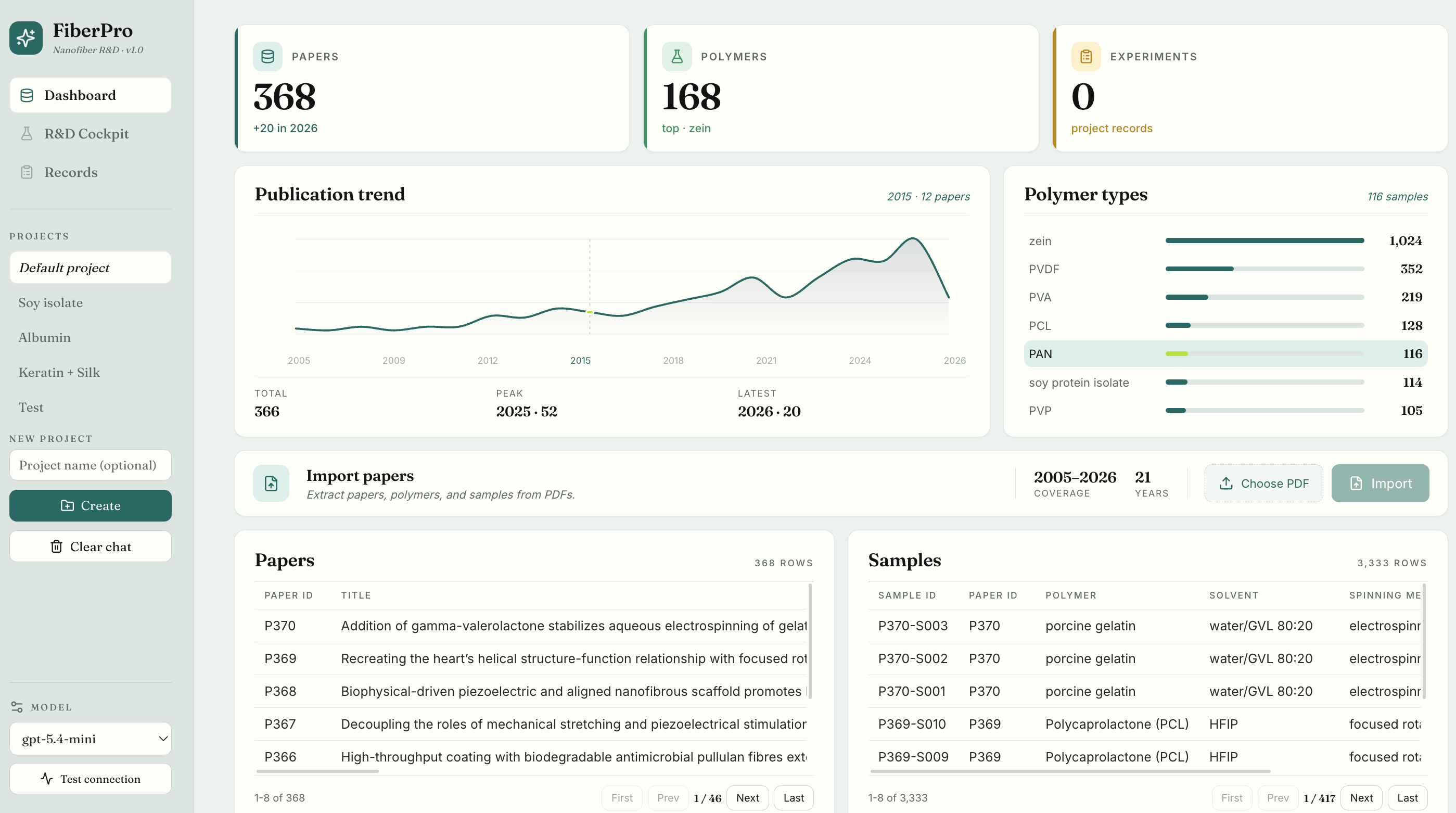

**Fig. S2. Dashboard of the FiberPro project database.** The dashboard summarizes project-specific knowledge, including literature coverage, polymer statistics, publication trends, and extracted experimental records. Users can import publications, browse structured paper- and sample-level databases, and track the expanding knowledge base to support materials design and experimental planning.

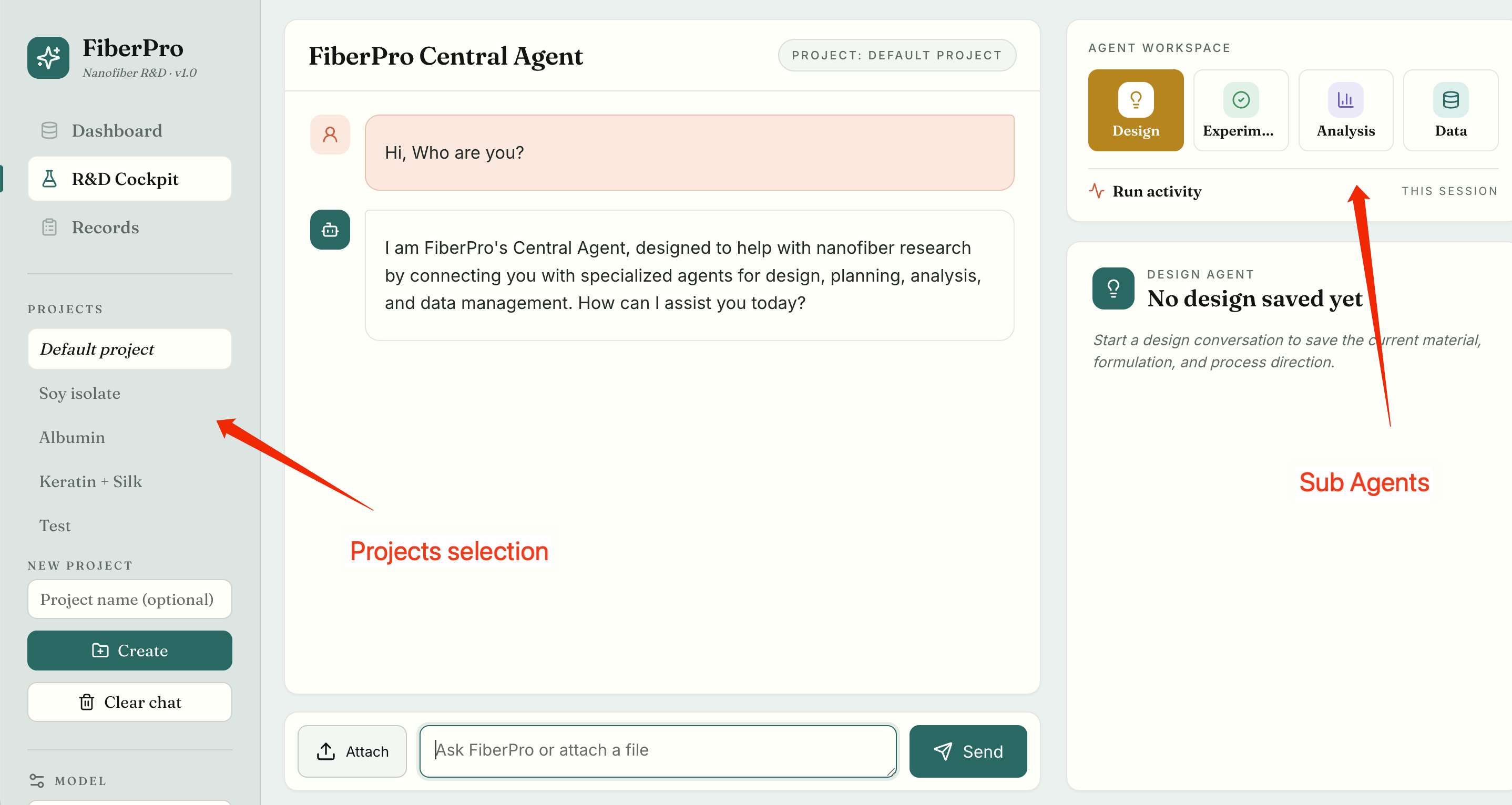

**Fig. S3. Graphical user interface of FiberPro 1.0.** The left sidebar enables project creation and selection while preserving project-specific context. The central workspace provides the primary interface for interaction with the Central Agent. The right-hand panel displays the available specialist agents and current run activity.

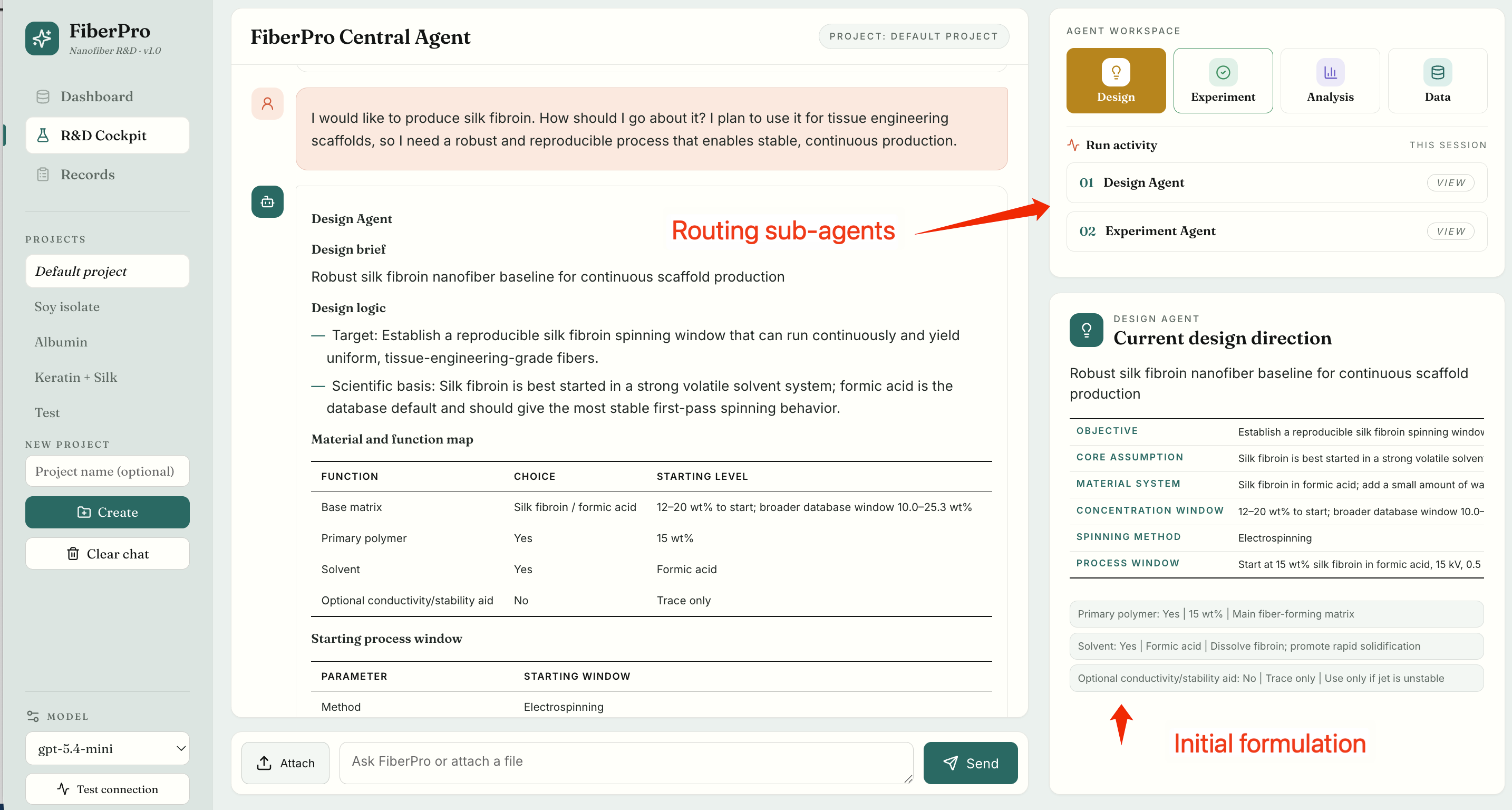

**Fig. S4. Example of a design workflow in FiberPro 1.0.** In response to a request for a robust silk fibroin formulation for continuous scaffold fabrication, the Central Agent routes the task to the Design Agent. The Design Agent retrieves project-relevant knowledge and proposes an initial formulation, together with a preliminary processing window and the corresponding design rationale. The resulting recommendation is displayed in the project workspace to support experimental planning and subsequent refinement.

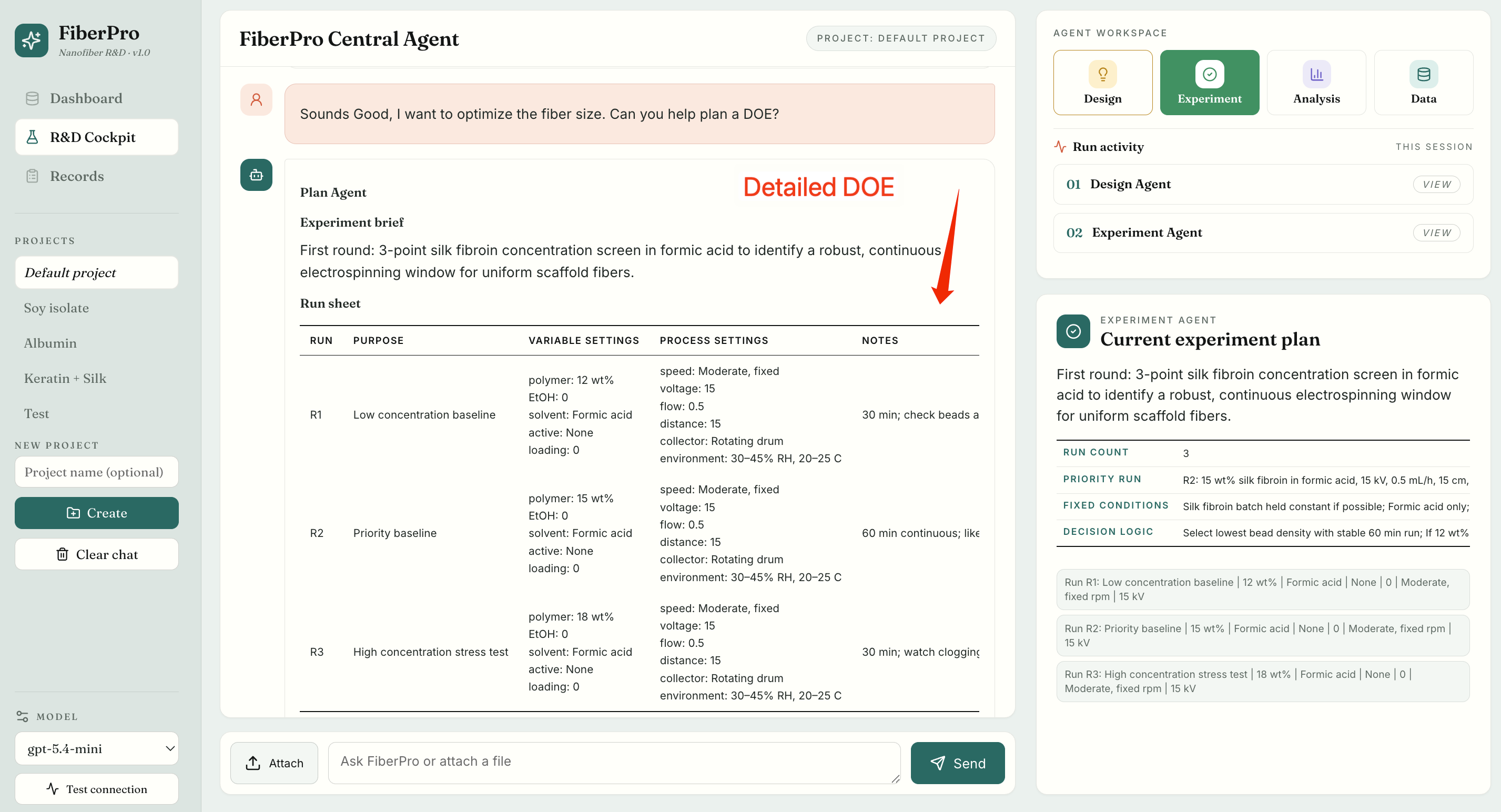

**Fig. S5.** Example conversion of an initial formulation into an experimental plan. Using the formulation proposed by the Design Agent, the Experiment Agent generates a structured design-of-experiments (DOE) plan for fibre-size optimisation. The plan defines which conditions should be varied, which should remain fixed, and what evidence is required before advancing to the next round. It is stored in the project memory to support reproducible execution and iterative optimisation.

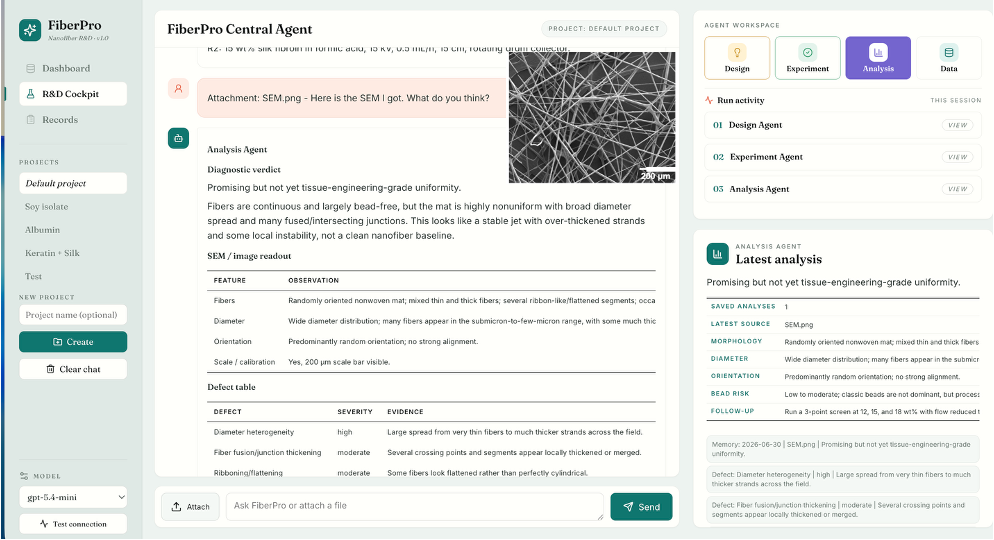

**Fig. S6. Example analysis workflow in FiberPro.** An SEM image uploaded to the workspace is assessed by the Analysis Agent, which provides a morphology-level assessment. The agent identifies observed defects, evaluates their likely causes, and recommends a targeted follow-up experiment. The resulting analysis is stored in project memory to inform subsequent materials design and experimental decisions.

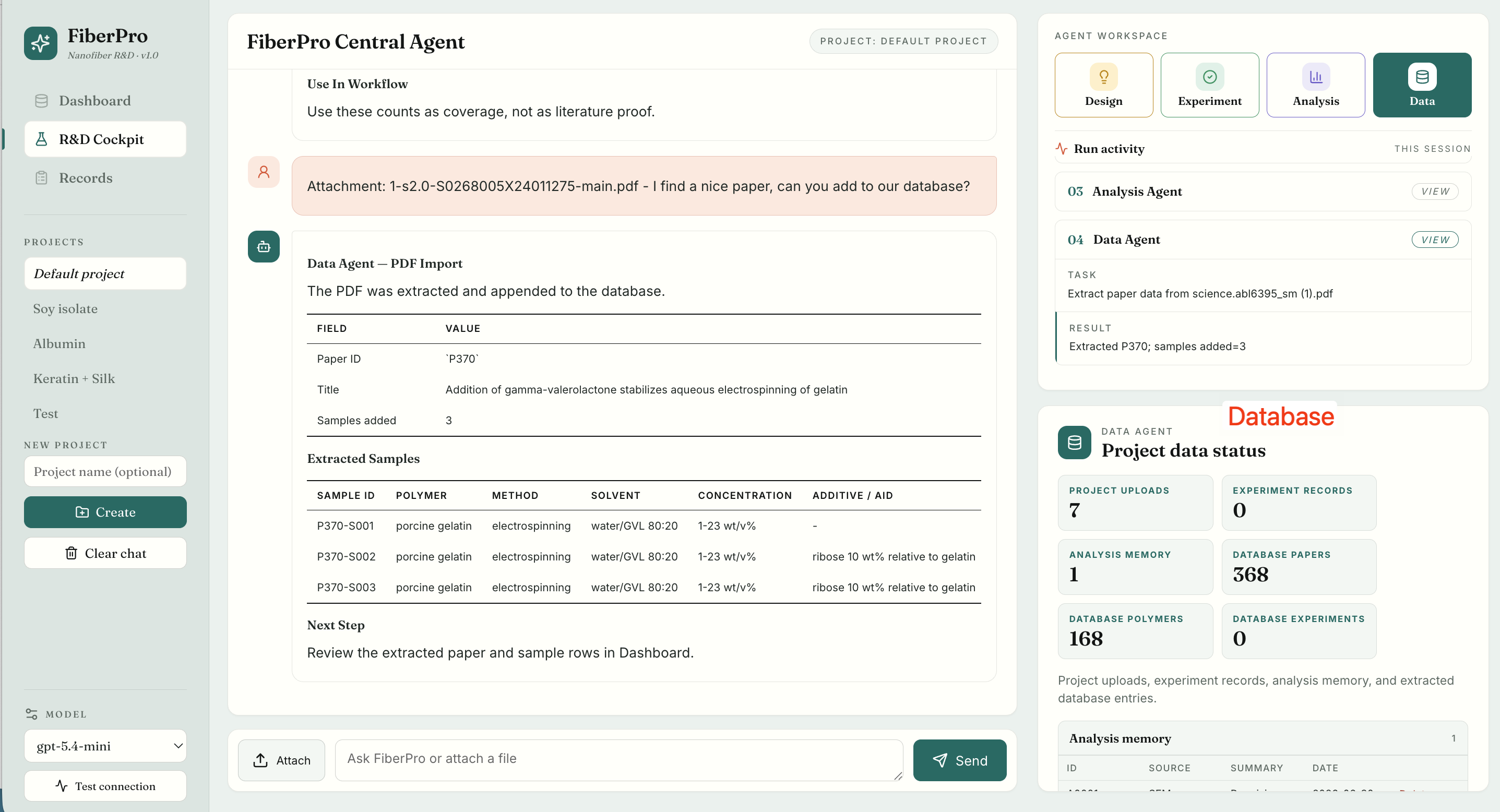

**Fig. S7. Example literature-to-database workflow in FiberPro**. A research article uploaded as a PDF is parsed by the Data Agent to extract article metadata and sample-level experimental records. The extracted information is incorporated into the project database, where the resulting structured records and project status can be reviewed. These records support subsequent knowledge retrieval for formulation design and experimental planning.

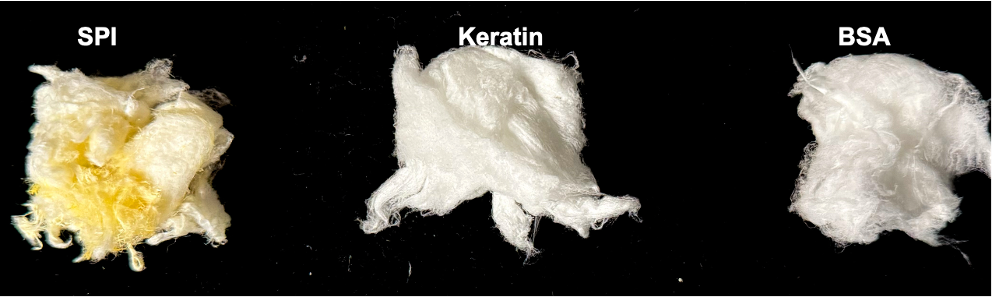

**Fig. S8. Representative photographs of protein fibers fabricated by FRJS from soy protein isolate (SPI), keratin, and bovine serum albumin (BSA).** The images show the macroscopic appearance of the resulting fiber samples produced under the feasible formulation conditions summarized in Table S1.

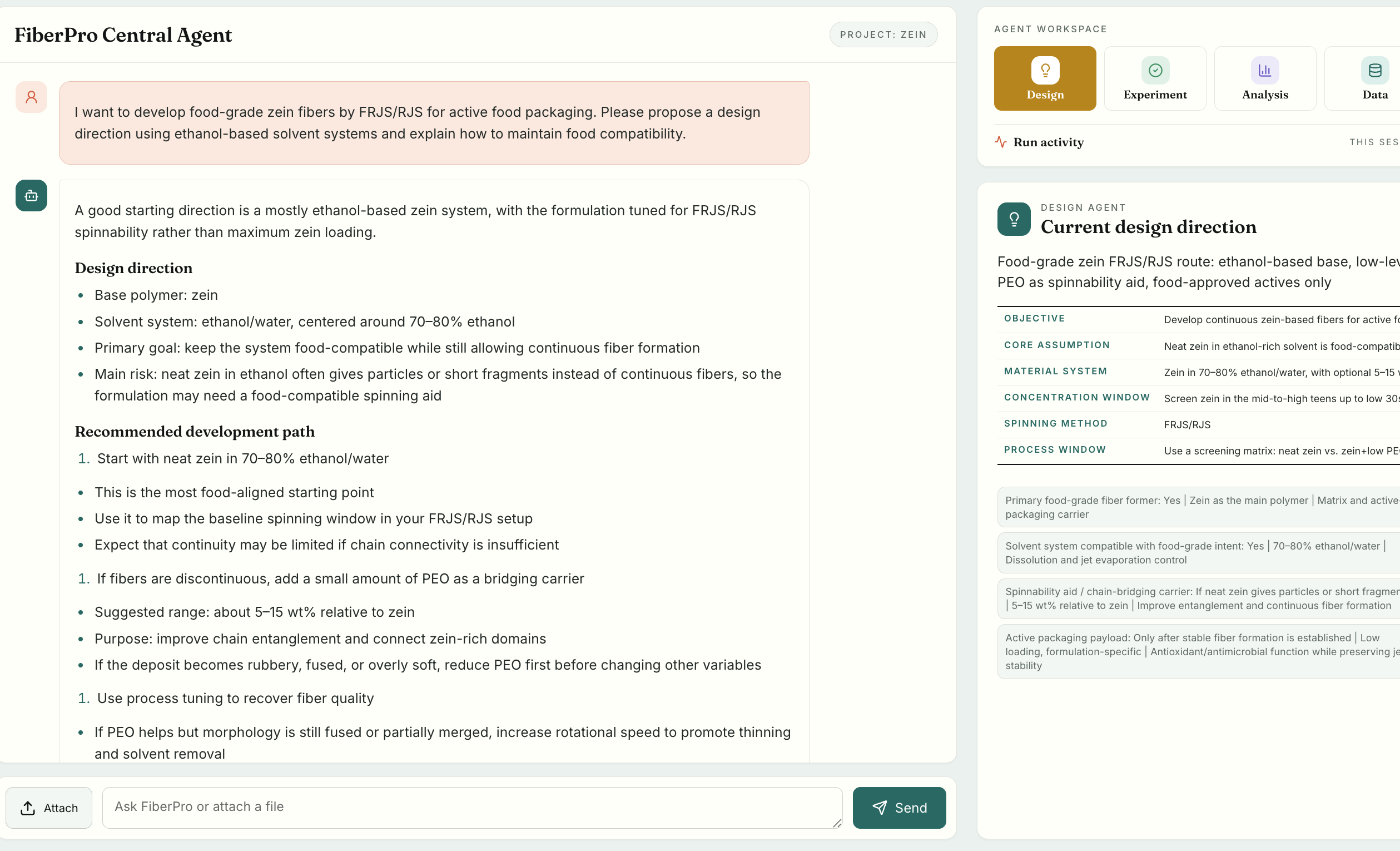

**Fig. S9. FiberPro-guided design of a zein-based fiber formulation for active food packaging.** The Central Agent proposed an ethanol/water-based zein formulation for focused rotary jet spinning (FRJS), prioritizing neat zein as the initial food-compatible system and providing a staged formulation strategy for subsequent optimization if continuous fiber formation was not achieved.

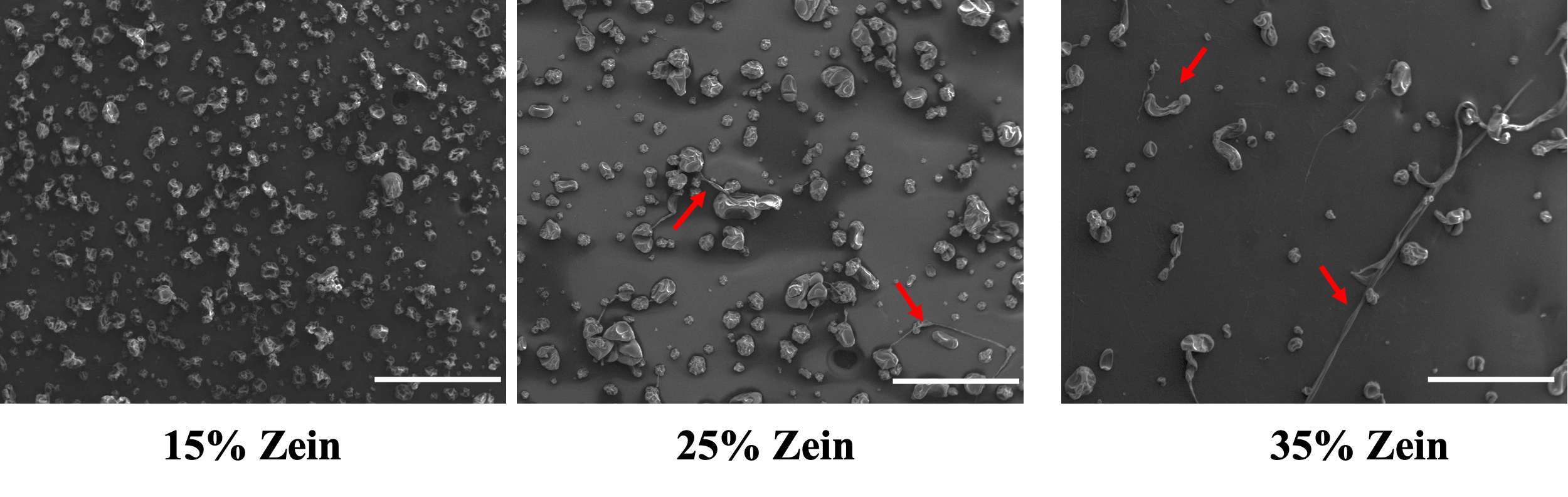

**Fig. S10. SEM images of products obtained from attempted FRJS of pure zein solutions.** The resulting structures consist predominantly of beads and aggregates, with occasional partially stretched jet segments indicated by red arrows. Scale bar, 500 μm.

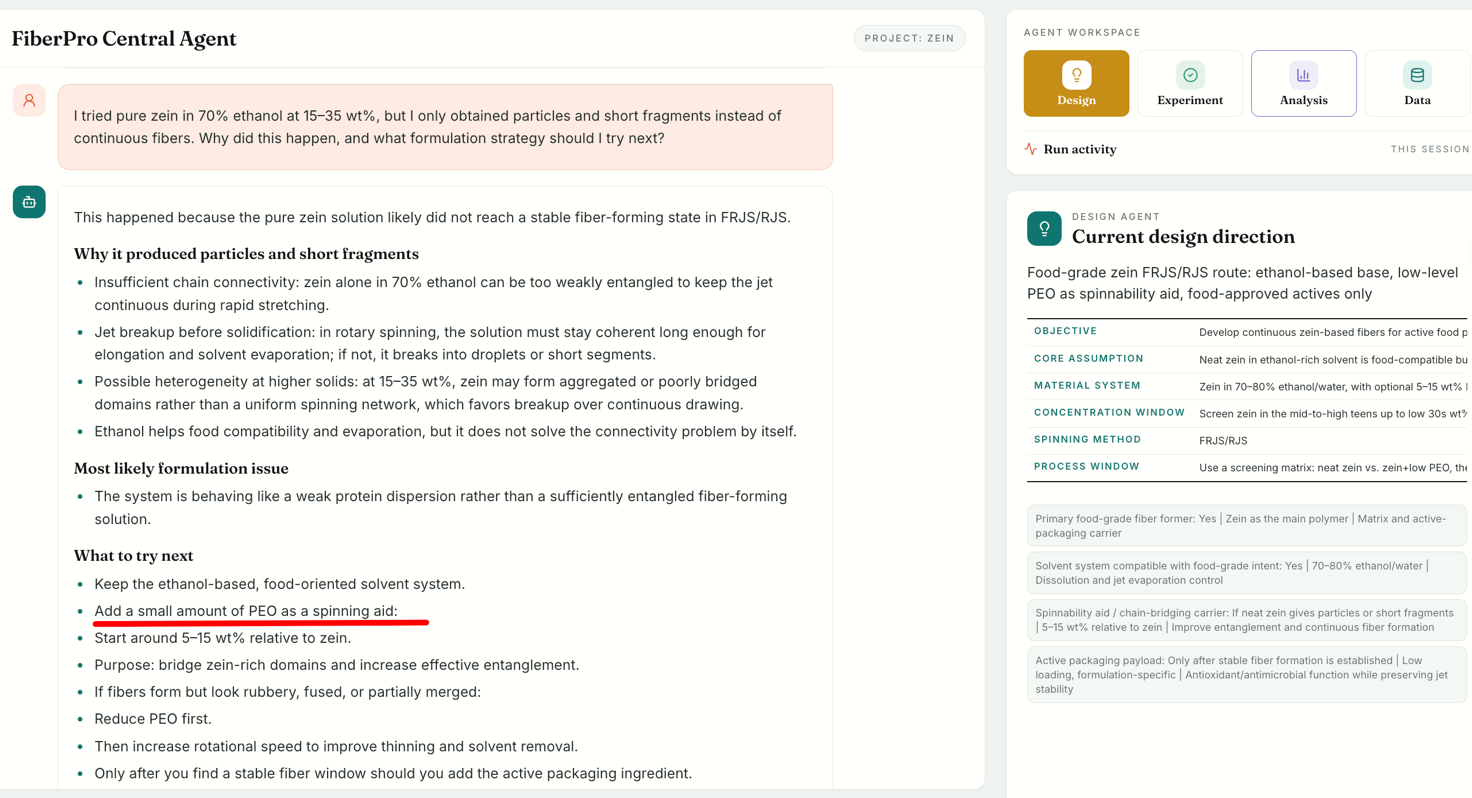

**Fig. S11. FiberPro diagnosis following unsuccessful spinning of pure zein.** After pure zein in 70% ethanol produced predominantly particles and short jet fragments, FiberPro interpreted the failure as insufficient chain entanglement and continuity of the fiber-forming network. The system recommended introducing a low concentration of PEO as a spinning aid to improve intermolecular connectivity and jet continuity.

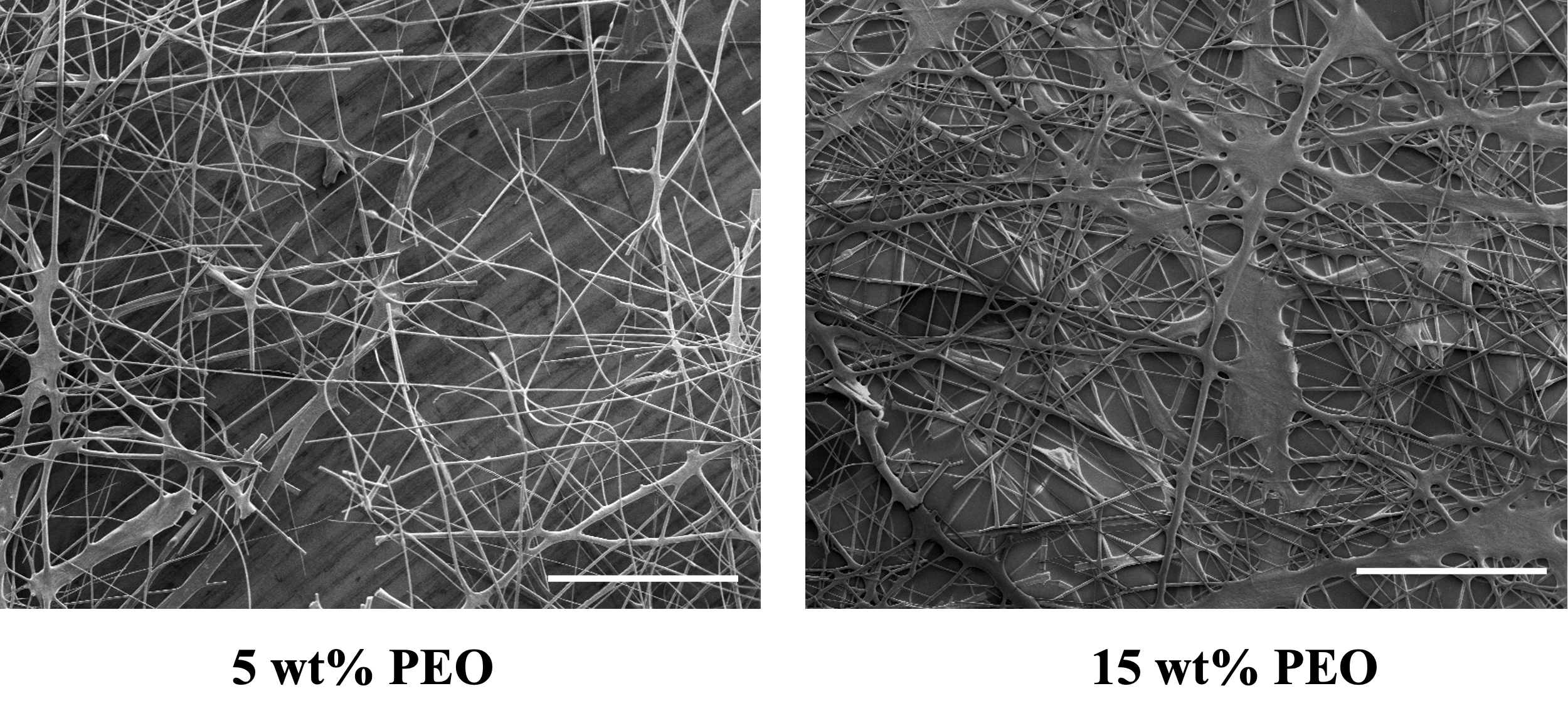

**Fig. S12. SEM images of zein/PEO fibers produced at different PEO loadings.** Left, 5 wt% PEO relative to zein, showing beaded fibers with intermittent membrane-like defects. Right, 15 wt% PEO relative to zein, showing extensively fused, ribbon-like structures with film-like junctions. Scale bar, 500 μm.

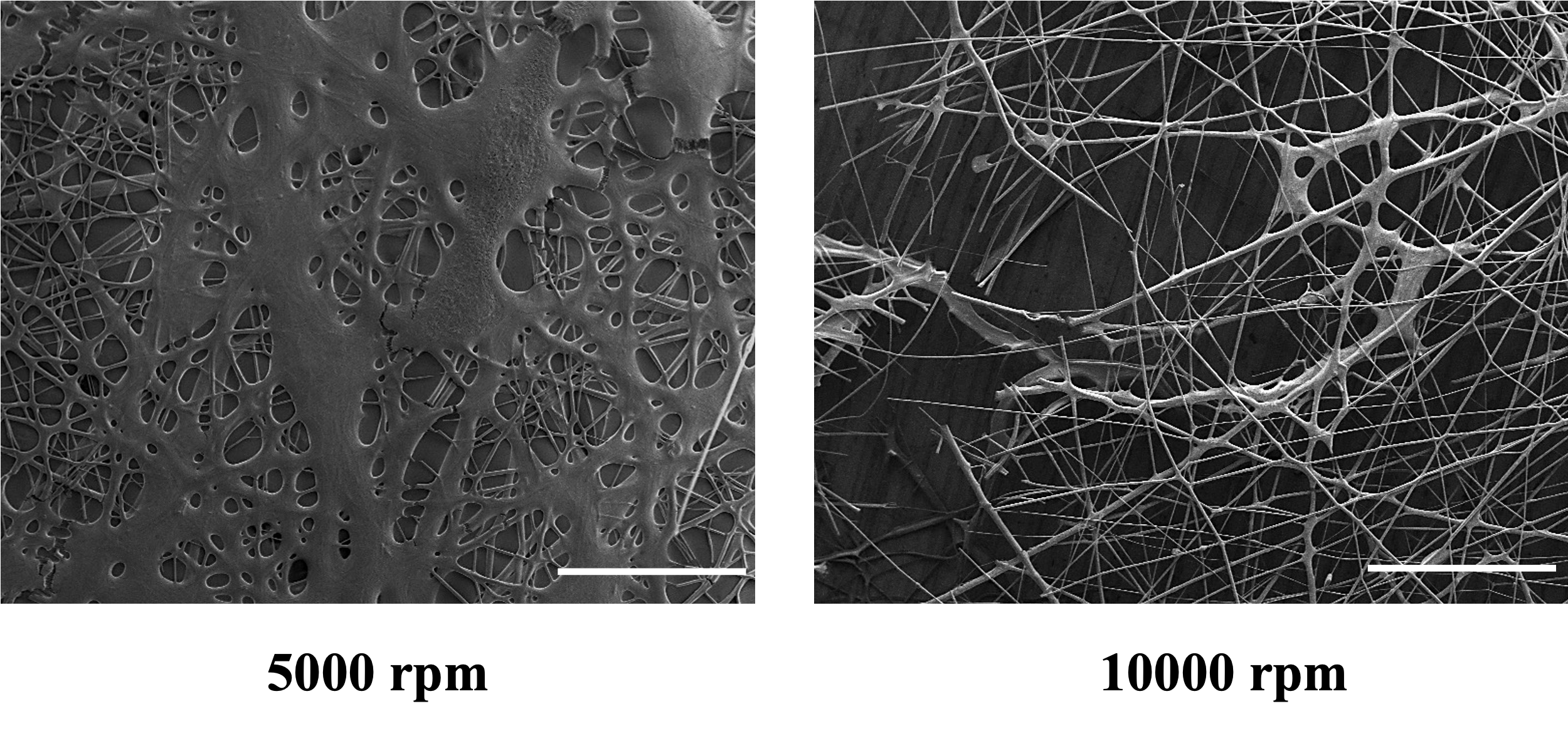

**Fig. S13. Effect of rotational speed on the morphology of zein/PEO fibers.** Left, fibers collected at 5,000 rpm, showing thick, highly fused networks with irregular morphology. Right, fibers collected at 10,000 rpm, showing thinner and more continuous fiber strands with reduced fusion. Scale bar, 500 μm.

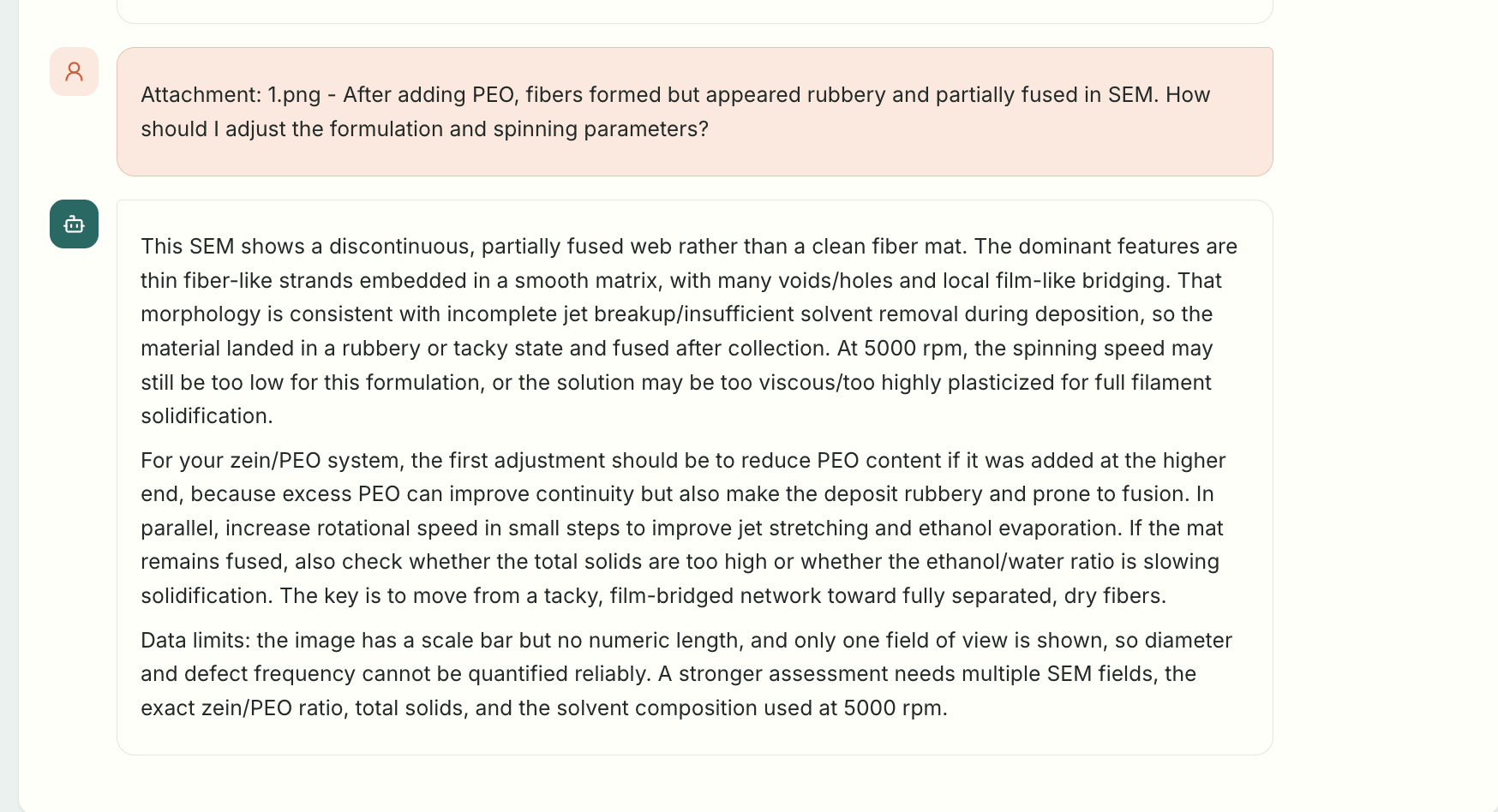

**Fig. S14. FiberPro analysis of fused zein/PEO fibers.** After PEO addition restored continuous fiber formation, SEM imaging revealed extensive fiber fusion and flattened or film-like junctions. FiberPro interpreted these features as being consistent with incomplete solvent removal and/or excessive plasticization associated with PEO and recommended reducing the PEO content while increasing rotational speed to enhance jet stretching and solvent evaporation.

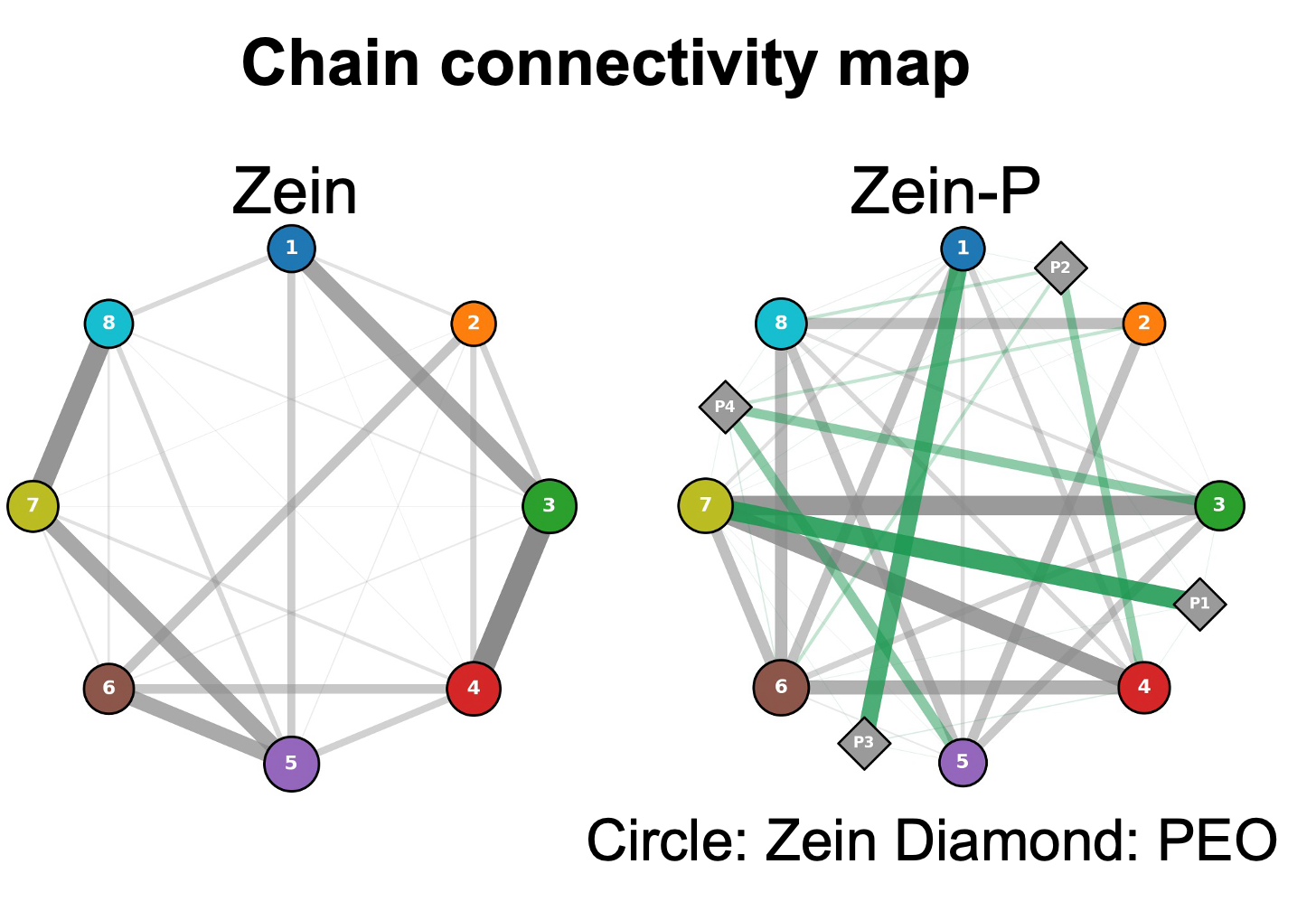

**Fig. S15.** Inter-chain contact graph at 10:30 for the zein (left) and zein + PEO (right) systems. Circles denote the eight zein chains and diamonds the four PEO chains; grey edges denote zein–zein contacts and green edges PEO–zein contacts, with edge width and opacity proportional to the number of atomic contacts (< 0.6 nm) averaged over the final 10 frames

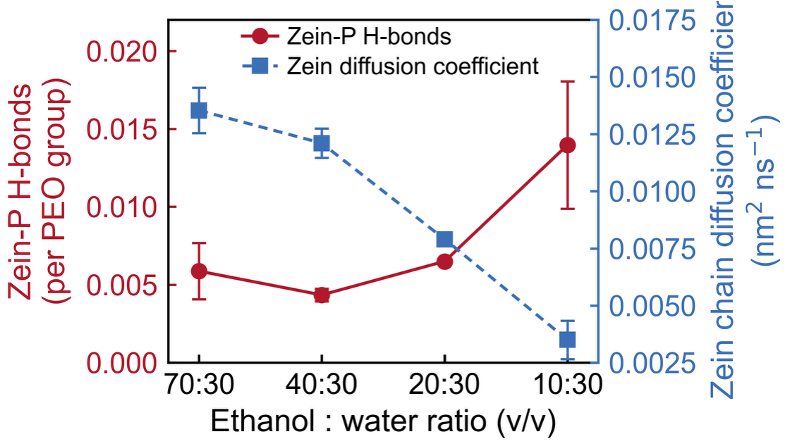

**Fig. S16.** Zein–PEO hydrogen bonds per PEO chain (red, left axis) and zein-chain diffusion coefficient (blue, right axis) as a function of ethanol:water ratio. Data are mean ± s.d. of three independent simulations.

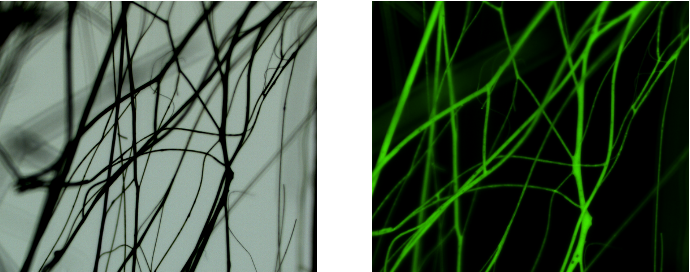

**Fig. S17. Bright-field (left) and fluorescence (right, GFP channel) micrographs of zein–curcumin (zein–CUR) fibers.** The fluorescence signal is distributed continuously along the fibers, indicating a relatively uniform distribution of curcumin within the fibrous structure.

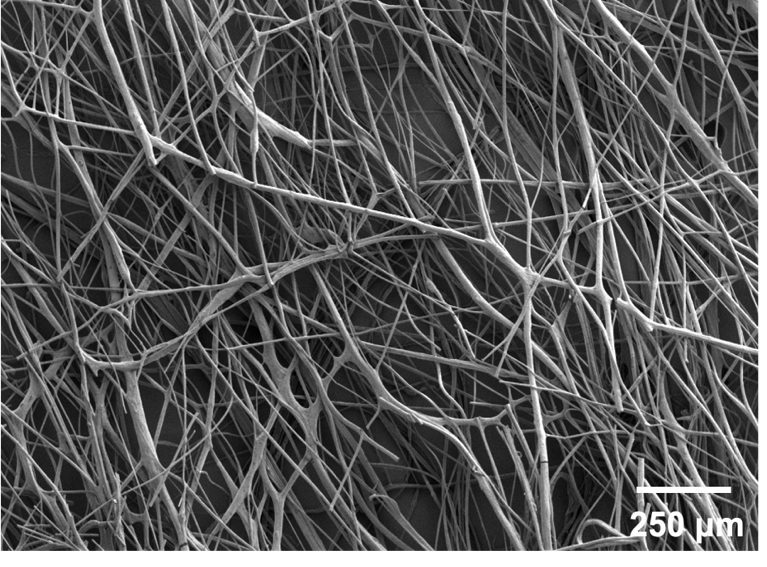

**Fig. S18.** Scanning electron micrograph of curcumin-loaded zein–PEO (Zein-PC) fibers, showing an interconnected fibrous network. Scale bar, 250 μm.

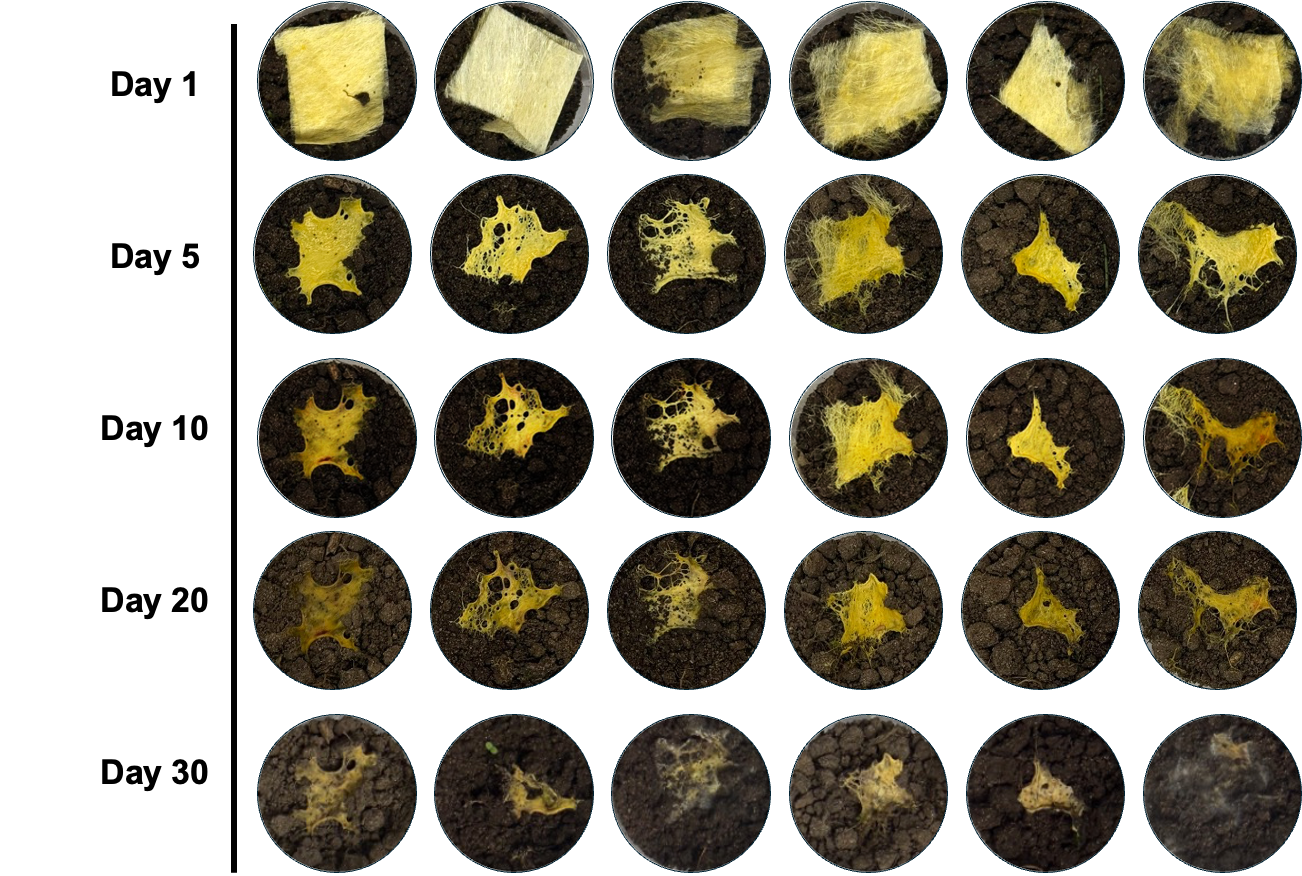

**Fig. S19. Macroscopic degradation of Zein-PC fibers over 30 days.** Representative photographs show the progressive loss of structural integrity and fragmentation of the fibrous samples during the degradation period.

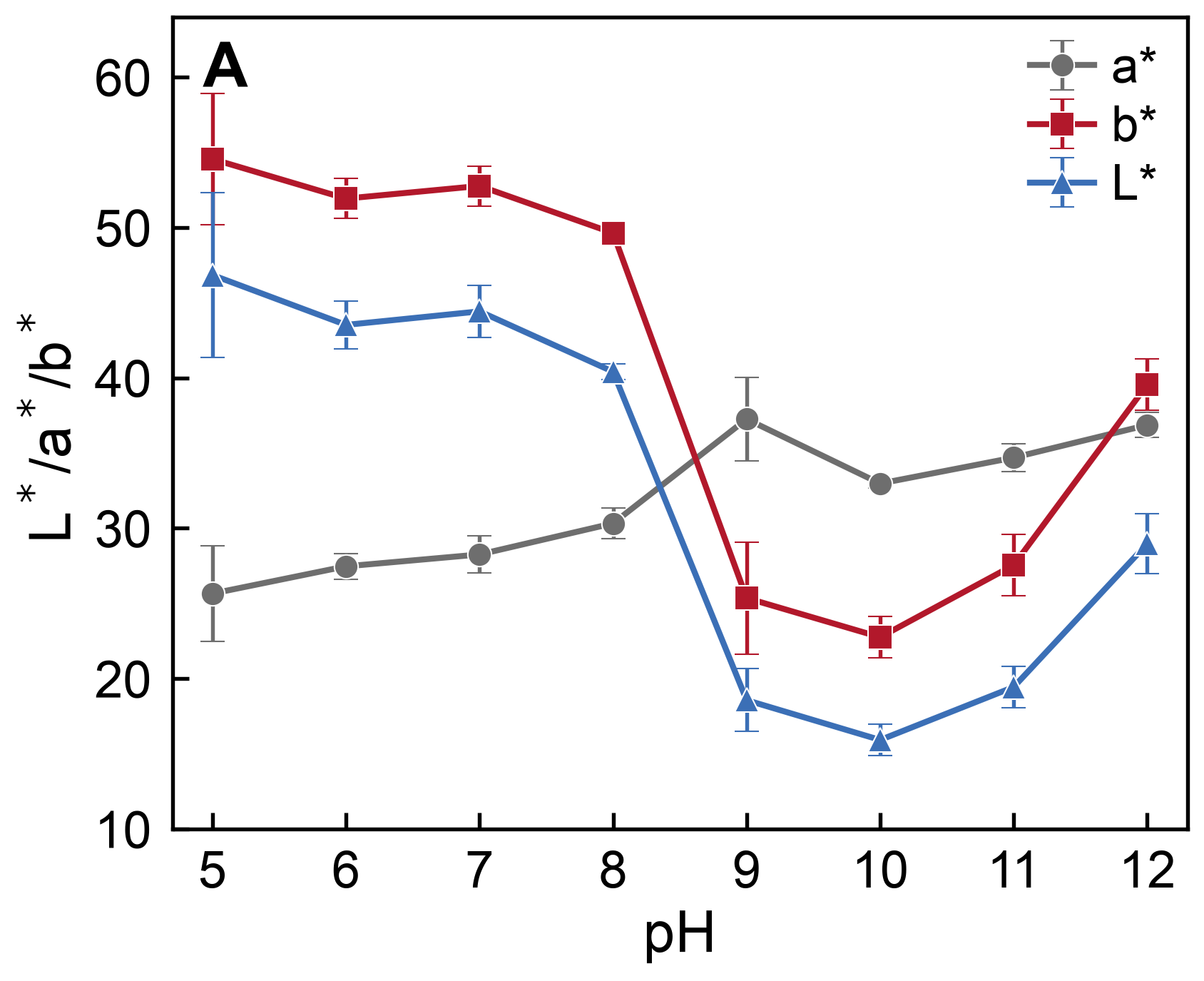

**Fig. S20. CIELAB color parameters of Zein-PC fibers as a function of pH.** The color coordinates (L*, a*, and b*) exhibit distinct pH-dependent changes, including a pronounced decrease in b* and an increase in a* under alkaline conditions, consistent with the characteristic yellow-to-red color transition of curcumin. Error bars represent SD (n = 3).

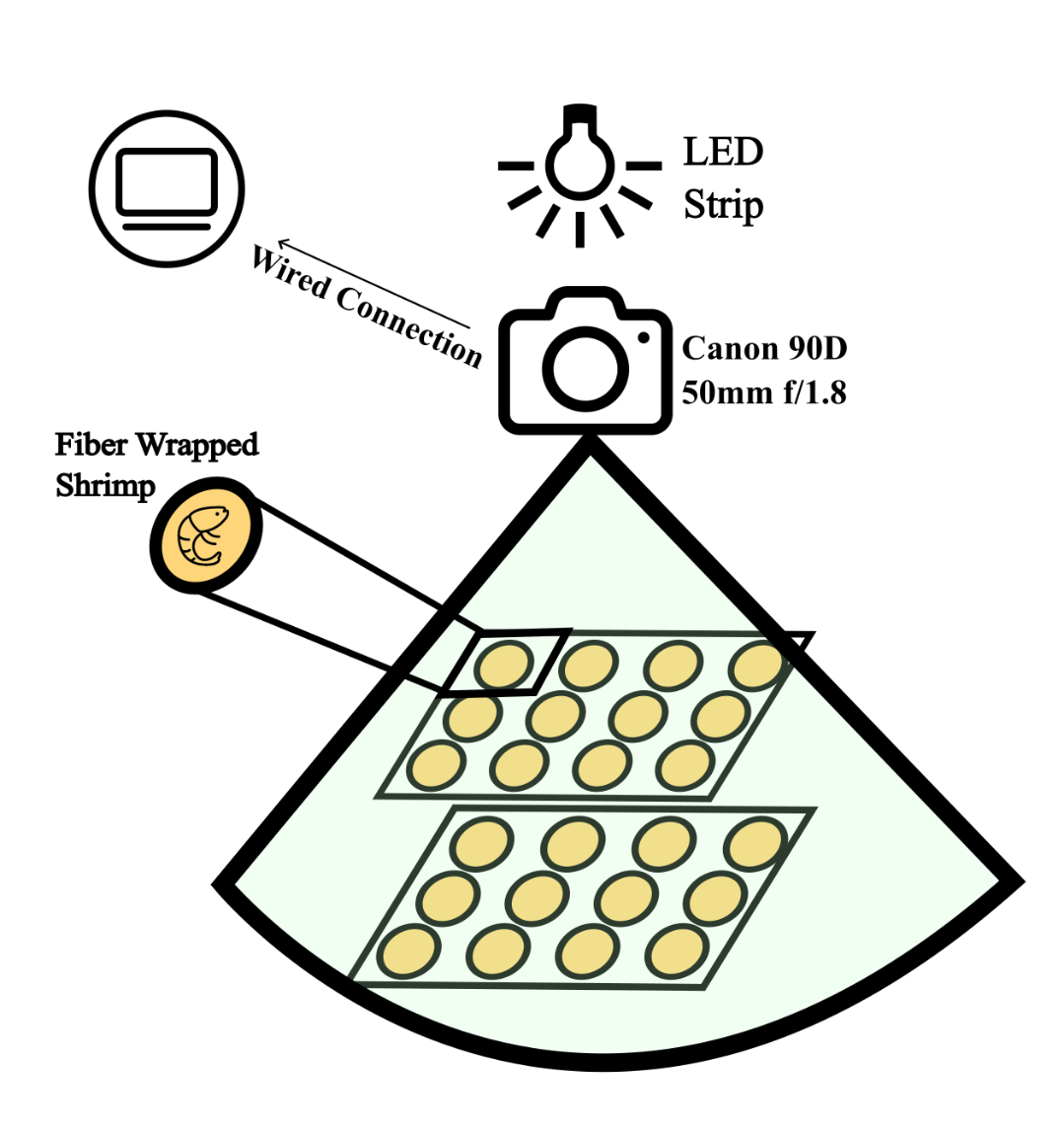

**Fig. S21. Schematic of the imaging setup used for visual and colorimetric analysis.** The system comprises an LED light source, a digital camera (Canon EOS 90D with a 50-mm f/1.8 lens), and a computer connected through a wired interface for image acquisition and camera control. Fiber-wrapped shrimp samples were positioned within a fixed field of view and imaged under controlled illumination to ensure consistent image acquisition.

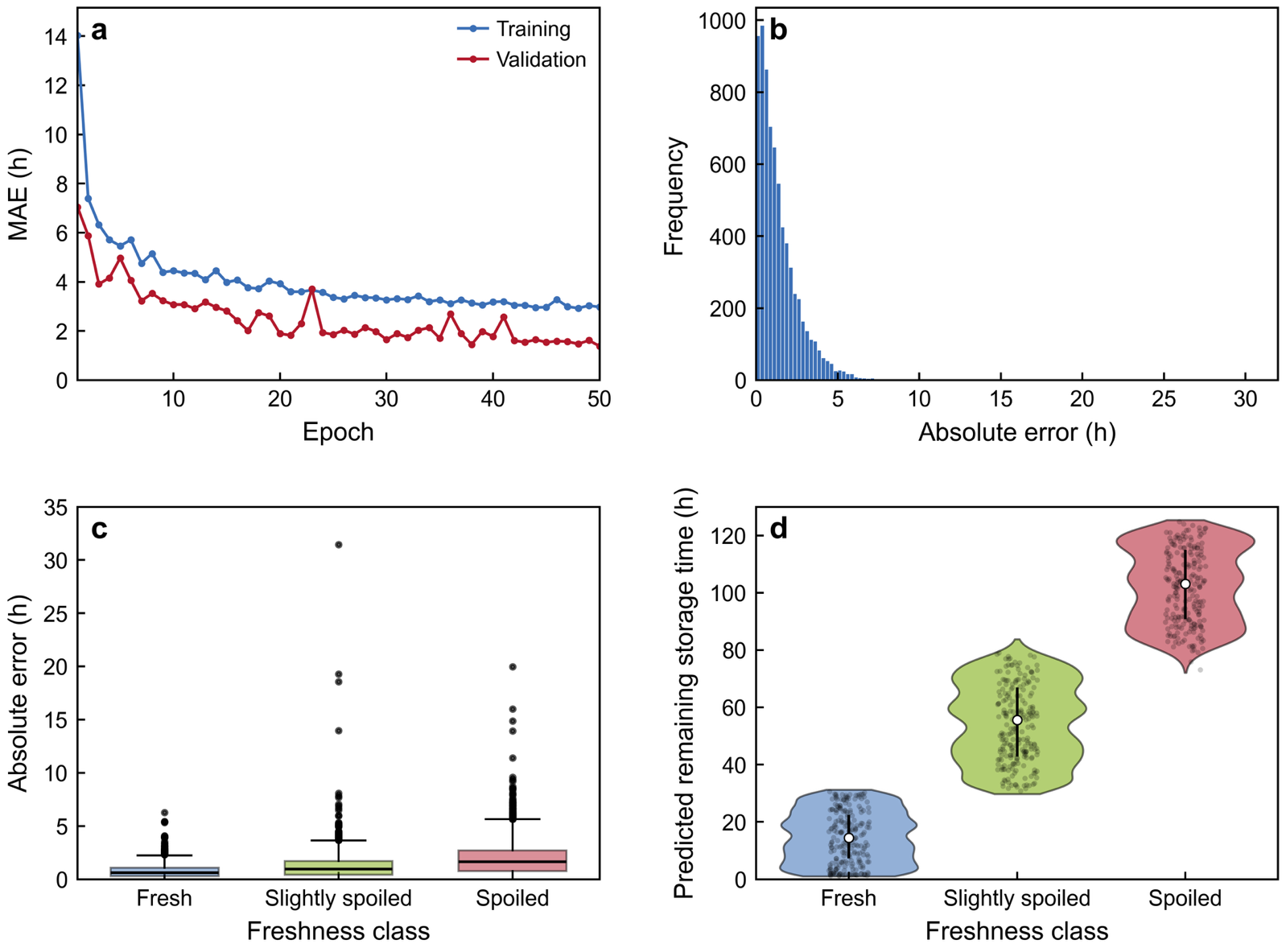

**Fig. S22. Training dynamics and regression diagnostics for the remaining shelf-life prediction model.** (A) Training and validation mean absolute error (MAE) over 50 epochs for the remaining shelf-life regression branch. (B) Distribution of absolute prediction errors across the test set. (C) Absolute prediction errors stratified by freshness class. Boxes indicate the interquartile range, center lines indicate medians, whiskers extend to 1.5 × the interquartile range, and points denote outliers. (D) Distributions of model-predicted remaining shelf life for fresh, slightly spoiled, and spoiled samples. Violin plots show kernel-density estimates; white circles and vertical bars indicate medians and interquartile ranges, respectively. Jittered black points represent randomly subsampled test-set predictions for visualization.

| **Protein** | **Round** | **Researcher-FiberPro dialogue and decision path** | **Result** |
| --- | --- | --- | --- |
| Soy protein isolate | **1** | **Researcher:** SPI does not dissolve into a homogeneous spinning solution.  **FiberPro:** Use alkaline dissolution and reassess solution homogeneity before FRJS. | Homogeneous alkaline solution obtained; fibres not yet collectable. |
|  | **2** | **Researcher:** The alkaline solution remains unsuitable for continuous fibre collection.  **FiberPro:** Introduce PVA to increase chain entanglement and validate the PVA-assisted formulation. | PVA-assisted condition selected for final validation. |
|  | **3** | **Researcher:** The PVA-assisted formulation yields continuous, collectable fibres.  **FiberPro:** Accept this condition as feasible and proceed to morphological characterisation. | First successful, collectable-fibre condition. |
| Keratin | **1** | **Researcher:** A high-keratin formulation is required without losing spinnability.  **FiberPro:** Use a formic acid/PVP formulation and verify fibre collection at high keratin content. | Stable, collectable fibres obtained at high keratin content. |
| Bovine serum albumin (BSA) | **1** | **Researcher:** An aqueous route is preferred over a conventional organic-solvent system.  **FiberPro:** Start with an aqueous glycerol/pullulan formulation to support fibre formation. | Aqueous formulation selected for validation. |
|  | **2** | **Researcher:** The aqueous formulation yields collectable fibres.  **FiberPro:** Accept this condition as feasible and retain it for further optimisation. | Collectable fibres obtained from the aqueous formulation. |

Table S1. Representative FiberPro dialogue records for iterative FRJS formulation development.
